# High-coverage DNA sequence and modification profiling of targeted genomic elements using Nanopore-based Cas12a Targeted Ligation and Enrichment Sequencing (nCasTLES)

**DOI:** 10.64898/2026.08.25.747114

**Authors:** Milo Vantine, Kensei Kishimoto, Brendan A. Pacheco, William A. Flavahan

## Abstract

Third-generation sequencing technologies, such as nanopore sequencing, enable long-read sequencing and direct characterization of nucleic acid modifications at low cost^1,2^. However, nanopore sequencing is limited by low throughput, necessitating targeted sequencing for interrogation of specific genomic elements. The current standard is nanopore Cas9-targeted sequencing (nCATS), which utilizes blunt-end cleavage of dephosphorylated DNA to render targeted DNA sites as the only ligation-capable ends for sequencing adapter addition^3^. nCATS significantly improves on-target sequencing yield but suffers from lower total sequencing output and faster flow cell degradation, resulting in an increased cost per sequencing due to inert DNA. Here, we present a modified approach, based on creating predictable base overhangs with Cas12a/Cpf1 as ligation substrates for biotinylated oligos followed by bead enrichment, termed nanopore Cas-12a Targeted Ligation-Enrichment Sequencing, or nCasTLES. nCasTLES removes off-target DNA via bead washes rather than rendering it inert. Removal of the inert off-target DNA allows nCasTLES libraries to be pooled with other sequencing libraries in a single sequencing run to achieve equivalent on-target DNA sequencing as nCATs while improving overall yield of useful data and decreasing the speed of flow cell degradation. We demonstrate the power of nCasTLES to characterize methylation dynamics at a frequently-methylated gene promoter. We also directed the Cas12a cleavage to an integrated lentiviral vector, allowing us to assess clonality of a transfected population and interrogate the integration state and transgene effects in selected clones. Finally, we demonstrate the utility of nCasTLES’ increased flow cell throughput by spike-in of nCasTLES libraries to WGS libraries to also characterize genetic and modified base information, such as clonal copy number variation analysis or BrdU incorporation, alongside the targeted sequencing. This approach will enable highly focused genomic interrogation in combination with full throughput of off-target reads.

## Introduction

Targeted enrichment is an essential method for fully leveraging third-generation sequencing, allowing for interrogations requiring longer read lengths or native base modification information from samples with limited input and with reduced labor and sequencing costs^4–6^. The current standard approach is nanopore Cas9-targeted sequencing (nCATS)^3^. nCATS starts by dephosphorylating input DNA and then using Cas9 to create phosphorylated blunt-end cleavage sites at targeted regions in the DNA; A-tailing and adapter ligation thus only occurs at these targeted sites. Notably, the off-target DNA is simply rendered inert rather than removed. For Oxford Nanopore Technologies’ (ONT) R9 flow cell, the presence of inert DNA in the sequencing mix did not adversely affect the sequencing, but the R10 may be more sensitive to inert DNA and with the shift to R10, ONT no longer sells a Cas-based sequencing kit. Additionally, target DNA is not enriched, simply rendered preferentially sequenceable. As such, in nCATS off-target inert DNA remains present during sequencing, which disallows library mixing and contributes to rapid degradation of the flow cell, resulting in greatly diminished throughput compared to a standard run. One of the inherent limitations of nCATS is that Cas9 creates blunt ends, rendering isolation of targeted DNA unfeasible with nCATS. Recent work has attempted to enrich for nCATS-ligated DNA through preferential binding of the polyT sequence in the nanopore sequencing adaptor to a bead-bound polyA sequence, but a true target-specific enrichment method has yet to be developed^7^.

## Results

In nCasTLES, we use Cas12a/Cpf1, which leaves 4-5 base overhangs, allowing for increased target specificity and for DNA isolation, as in Cas12-Capture^8,9^. Cpf1 cleavage has been shown to cleave the target strand (TS) and non-target strand (NTS) at multiple locations, generating a variety of overhangs for a single target sequence^10^. By characterizing the site-specific overhangs left by Cas12a at each target site through amplicon sequencing, we create matching overhangs on biotinylated oligos and use streptavidin bead enrichment to isolate target DNA. Notably, our oligo design is structured such that the biotin is attached to the non-overhang strand, thus each individual site requires only a standard oligo to anneal to a single universal biotinylated oligo, reducing method cost and complexity. These biotinylated DNA adapters are next ligated only to the on-target Cas12a-generated overhangs, followed by optional endonuclease digest to reduce size of target DNA fragment to <10kb. Next, target DNA is bound to streptavidin beads and off-target DNA is washed off. Endonuclease digest is then used to cleave the biotin tag off the adapter and release the on-target DNA from the streptavidin beads. On-target DNA is then end-repaired and dA-tailed before ligation to nanopore sequencing adapters. The on-target DNA library can then be spiked-in to a WGS library, or any other DNA library of choice, and sequenced on a MinION flow cell (**Fig. 1a**).

**Figure 1.**
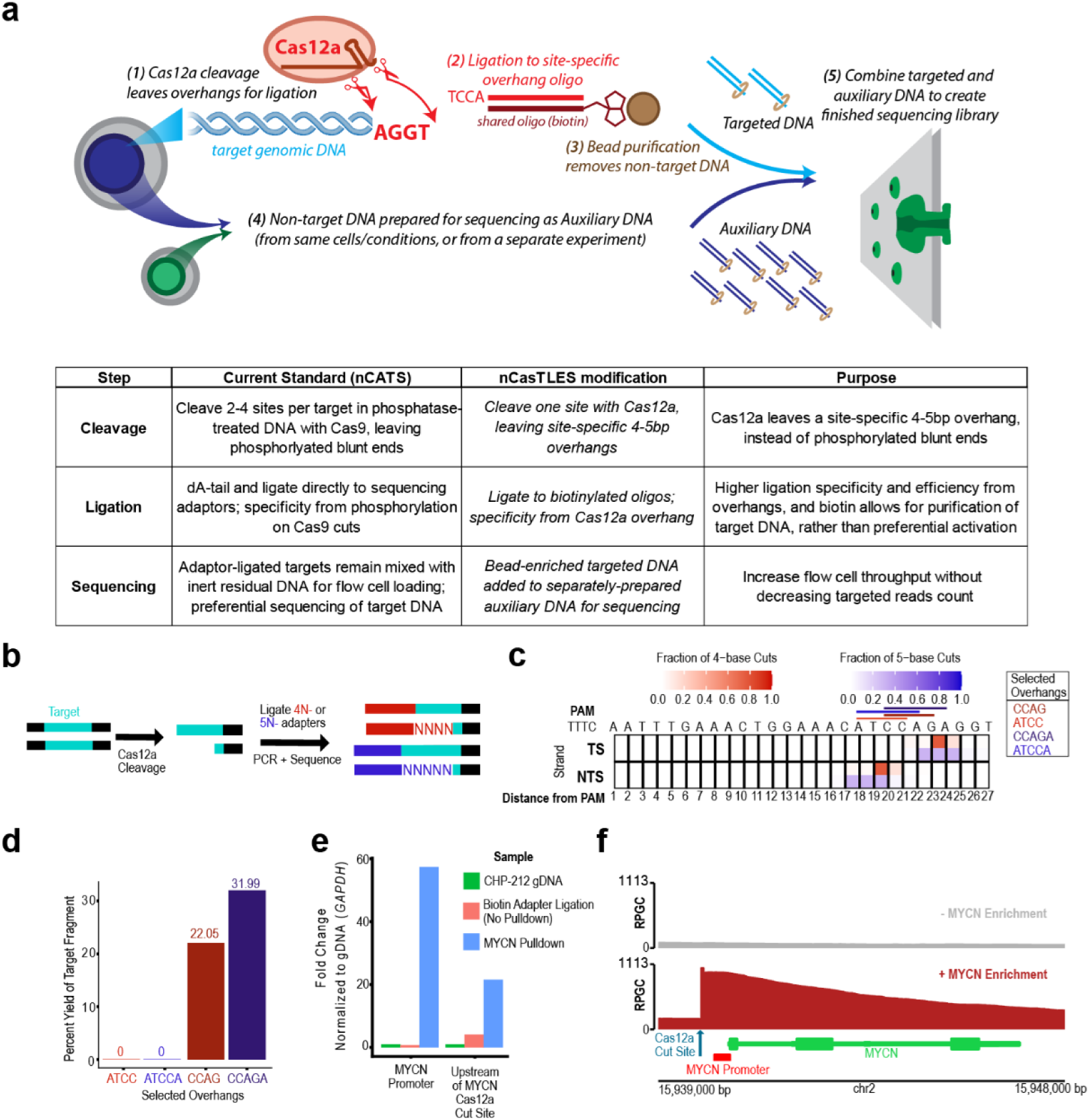
nCasTLEs overview, optimization, and validation. **(a)** Overview of nCasTLES. Targeted site is cleaved by a single Cas12a guide, leaving a predictable 4-5bp overhang. A site-specific oligo annealed to a common biotinylated oligo is ligated to the overhang. Captured fragments are then purified via streptavidin beads, and released from the bead by oligo cleavage. Separately, a second set of DNA is prepared for sequencing as the auxiliary DNA; this can be gDNA from the same cells, or a separate cell line, condition, or experimental setup. Both the enriched target DNA and the auxiliary DNA are combined to form the final sequencing library. Table presents differences between nCasTLES and the current standard approach, nCATS. (**b**) Cas12a site-specific overhangs are characterized via ligation of pooled 4- or 5-base overhang complementary oligos containing all possible overhangs. Successful ligation products are amplified via PCR and sequenced. (**c**) Plot depicts results of overhang analysis for *MYCN* Cas12a cleavage. Based on cleavage site frequency, four overhangs were selected for validation. (**d**) Bar plot displays validation pulldown of overhangs via qPCR. Of the four predicted overhangs, two successfully ligated to a Cas12a-cleaved *MYCN* DNA amplicon and enriched via streptavidin bead pulldown. (**e**) Bar plot displays successful enrichment of *MYCN* ecDNA via bead pulldown tested by qPCR. (**f**) Genomic enrichment tracks at *MYCN* demonstrate strong enrichment of *MYCN* via nCasTLES.

nCasTLES demonstrates improvements over current targeted nanopore-sequencing methods. nCasTLES achieves target enrichment with only one guide and one Cas12a cut per target, unlike nCATs, which requires dual Cas9 digest on either side of a locus of interest and up to 4 guides per target. Further, in nCATs target DNA is sequenced in the presence of abundant inert, off-target DNA, which cannot be sequenced. In contrast, nCasTLES depletes off-target DNA prior to sequencing and allows for sequencing of auxiliary DNA of choice, which improves overall flow cell health and increases the amount of usable data collected per sequencing run for each flow cell.

### Characterization of an oncogene-containing extrachromosomal DNA amplification

We first sought to deploy nCasTLES to understand a complex genetic driver alteration. CHP-212 cells contain a high copy number ecDNA composed of segments of chromosome 2, including the locus of the oncogene *MYCN*^11^. We sought to utilize this high copy-number target for our pilot nCasTLES sensitivity testing and optimization.

We designed a Cas12a guide targeted upstream of the MYCN promoter on the ecDNA using CRISPOR^12^ and confirmed efficient cleavage on a PCR template and isolated genomic DNA (**Sup. Fig. 1a,b**). We next profiled the staggered 5’ overhangs generated by Cas12a at the MYCN locus by digesting CHP-212 genomic DNA with the Cas12a-MYCN guide ribonucleoprotein (RNP), then ligating PCR adapters with either 4N- or 5N-variable overhangs (**Fig 1b**). PCR over the cleavage site was performed and the overhangs were identified with long-read amplicon sequencing (**Fig. 1c**). The top 4 overhangs detected in the amplicon sequencing data were selected and used to design site-specific DNA adapters. These adapters were tested via ligation and pulldown of a PCR product, and two successful hits were used to pull down CHP-212 genomic DNA, which was strongly enriched by qPCR (**Fig. 1d**., **Sup. Fig. 1c,d**).

To assess the performance of nCasTLES, we performed a pulldown of the MYCN ecDNA locus from CHP212 gDNA using our MYCN guide RNA. Comparison of on-target reads from the nCasTLES library to the same gDNA sequenced without enrichment revealed greatly improved coverage of the MYCN locus (**Fig. 1e**). Without enrichment, we captured 6.6 on-target reads per hour of sequencing. In contrast, with nCasTLES we captured 247.3 on-target reads per hour of sequencing. This ∼40-fold increase in on-target read capture per hour is achieved with just a single guide. Over the 34kb target region spanning from the Cas12a cut site to the downstream SbfI-HF cut site, the median coverage was 69X without enrichment and increased to 190X with nCasTLES (**Fig. 1f**). Taken together, this data shows that nCasTLES improves on-target capture and sequencing at a locus of interest over standard nanopore sequencing alone.

### Isolation and characterization of a non-amplified disease-relevant locus

We next sought to deploy nCasTLES to profile a non-amplified disease-relevant genomic target. The DNA repair enzyme methylguanine methyltransferase (MGMT) can repair alkylating chemotherapy damage, but is frequently inactivated by promoter methylation in glioblastoma (GBM)^12^. We have previously shown, via nCATS, that the GBM cell line T98G contains cells with both methylated and unmethylated promoters, which affects responses to temozolomide^14^. We designed a Cas12a guide targeted downstream of the MGMT promoter (**Fig. 2a**) using CRISPOR^12^ and confirmed efficient cleavage *in vitro*. We profiled the overhangs left by Cas12a at the MGMT cleavage site via amplicon sequencing and designed six overhangs; four of these demonstrated successful genomic DNA-pulldown by qPCR (**Fig. 2c, Sup. Fig. 2**).

**Figure 2.**
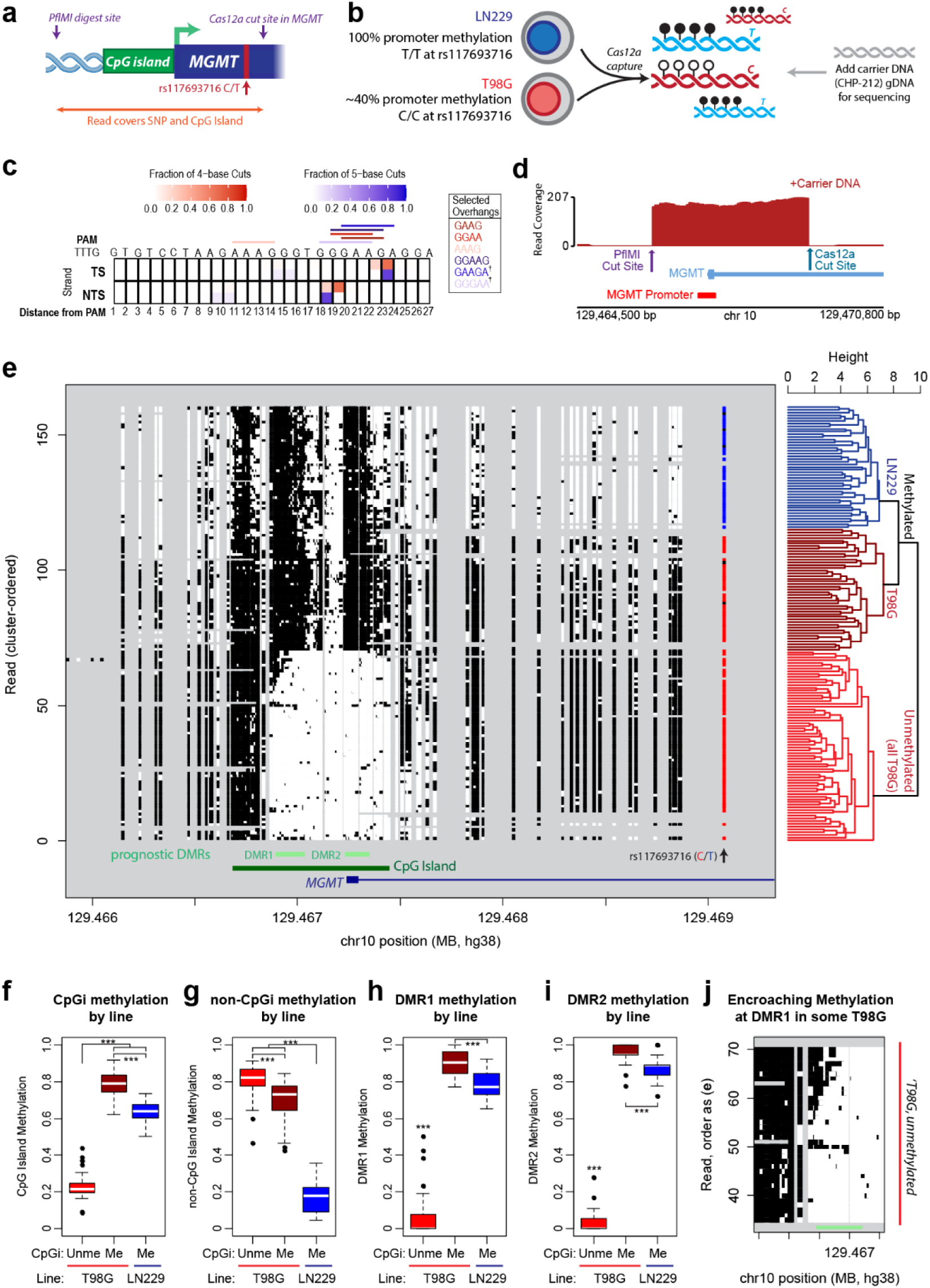
Characterization of a disease-relevant promoter methylation with nCasTLES. (**a**) Schematic depicts MGMT nCasTLES design strategy. A single Cas12a cut site within the MGMT gene will allow capture of the first exon of *MGMT*, the CpG Island at its promoter, and the SNP rs117693716. (**b**) Schematic depicts experimental setup. Genomic DNA from two glioblastoma lines, T98G and LN229, undergoes nCasTLES capture as described in (a) and is mixed and sequenced together. T98G and LN229 are each homozygous for opposite alleles at rs117693716, allowing for cell-line-of-origin determination for each sequenced molecule. nCasTLES-enriched GBM DNA is added to CHP-212 auxiliary DNA prepared by native barcoding kit for sequencing. (**c**) Heatmap depicts results of *MGMT* Cas12a overhang characterization. Six predicted overhangs were tested, resulting in four which were successfully able to enrich *MGMT* DNA (see Sup. Fig. 2, † indicates overhangs that did not validate). (**d**) Genomic traces demonstrate successful enrichment of *MGMT* promoter via nCasTLES. (**e**) Plot depicts nCasTLES *MGMT* promoter sequencing data. Dots depict CpGs, organized by read (rows) and position (columns), with white dots representing unmethylated CpGs and black representing methylated CpGs. Cell line SNP calls are indicated, with blue representing T calls (corresponding to LN229) and red representing C (T98G). Genomic elements, including the CpG island, prognostic differentially methylated regions (DMRs), and *MGMT* gene are indicated below. To the right, hierarchical clustering of reads by CpG methylation is presented, showing three clear clusters; first separating reads by CpG island methylation status, and then separating methylated reads by cell-line-of-origin. (**f-i**) Average methylation of CpGs in each read, separated by line and CpG island methylation status, are presented for CpGs within the CpG island (**f**), outside of the CpG island (**h**), or within each prognostic DMR (**i,j**). (**j**) Plot depicts subset of panel (e), focused on the reads classified as unmethylated T98G reads that displayed some methylation within DMR1.

An advantage of targeted nanopore sequencing approaches is the ability to directly profile read-level CpG methylation on native DNA without the need for bisulfite conversion. To demonstrate nCasTLES’ ability to profile diverse patterns of locus-specific CpG methylation, we performed nCasTLES on a mixture of genomic DNA from T98G cells and another GBM cell line, LN229, in which the promoters of *MGMT* are uniformly methylated (**Fig. 2b**). Notably, MGMT is expressed in LN229 cells, but not in T98G cells (**Sup. Fig. 3**). These captured MGMT promoters were then spiked into CHP-212 gDNA prepared with the native barcoding kit and run on the nanopore, which resulted in strong *MGMT* promoter coverage (**Fig. 2d**). We first performed hierarchical clustering on on-target MGMT promoter reads by methylation patterns, then assigned read origins (LN229 or T98G) by a cell line specific single nucleotide polymorphism (SNP) at rs117693716 (T/T in LN229, C/C in T98G) (**Fig. 2e**). As expected, hierarchical clustering revealed three main clusters of reads; the first separation was between reads with methylated and unmethylated CpG islands at the promoter, then within the methylated reads, two clear clusters emerged. Correlating these with the line-specific SNPs showed that these clusters corresponded to unmethylated T98G reads, methylated T98G reads, and methylated LN229 reads. Further analysis showed that the methylation patterns and levels within the methylated CpG islands differed between T98G and LN229 promoters (**Fig. 2e,f**), specifically the methylated T98G promoters had higher levels of methylation than the methylated LN229 promoters. Outside of the CpG island (i.e. upstream and intronic CpGs), T98G promoters were more similar to each other regardless of CpG island methylation state and uniformly displayed methylation, while LN229 promoters were uniformly unmethylated at these CpGs (**Fig. 2g**). Two prognostic differentially methylated regions (DMRs) within the CpG island have been reported^15,16^; we observed methylation at these locations to strongly correlate with each other and overall CpG island methylation, with several exceptions (**Fig. 2h,i**). Several reads, which were clustered and SNP-identified as unmethylated T98G promoters, displayed encroaching methylation spreading into DMR1 (**Fig. 2j**). The significance of this methylation is unclear; perhaps these reads represent a transient state, of either methylation spreading into an unmethylated promoter in the process of being silenced, or demethylation spreading outward in a reactivating previously methylated promoter. Additionally, we demonstrated that nCasTLES requires relatively low input DNA compared to nCATS: only ∼1ug of gDNA was required from each cell line for each MGMT nCasTLES run, contrasted with the ∼3ug recommended for the nCATS protocol. Altogether, we have demonstrated that we can utilize nCasTLES to simultaneously profile diverse patterns of CpG methylation and identify single nucleotide variants at high coverage without high input DNA requirements.

#### nCasTLES improves on-target coverage of the MGMT Locus with or without auxiliary DNA

During these experiments, we aimed to determine whether the addition to the nCasTLES library of a separate auxiliary WGS library affected the on-target read return. We ran one of the MGMT libraries alone and one added to a separately-prepared CHP-212 gDNA WGS library for sequencing. The rate of capture of on-target DNA sequences was much higher with the auxiliary library than without (Table 1). Furthermore, only 13.3% of the total reads sequenced in the nCasTLES run lacking auxiliary DNA mapped to hg38, with a large portion of total reads being short, unmappable “junk” DNA. In contrast, 91% of the total reads sequenced in the run with the WGS library mapped successfully to hg38. These data suggest that addition of nCasTLES libraries to an auxiliary library may increase on-target sequencing recovery while also providing useful additional data from sequencing the auxiliary DNA, improving both targeted and total flow cell data collection efficiency.

**Table 1.**
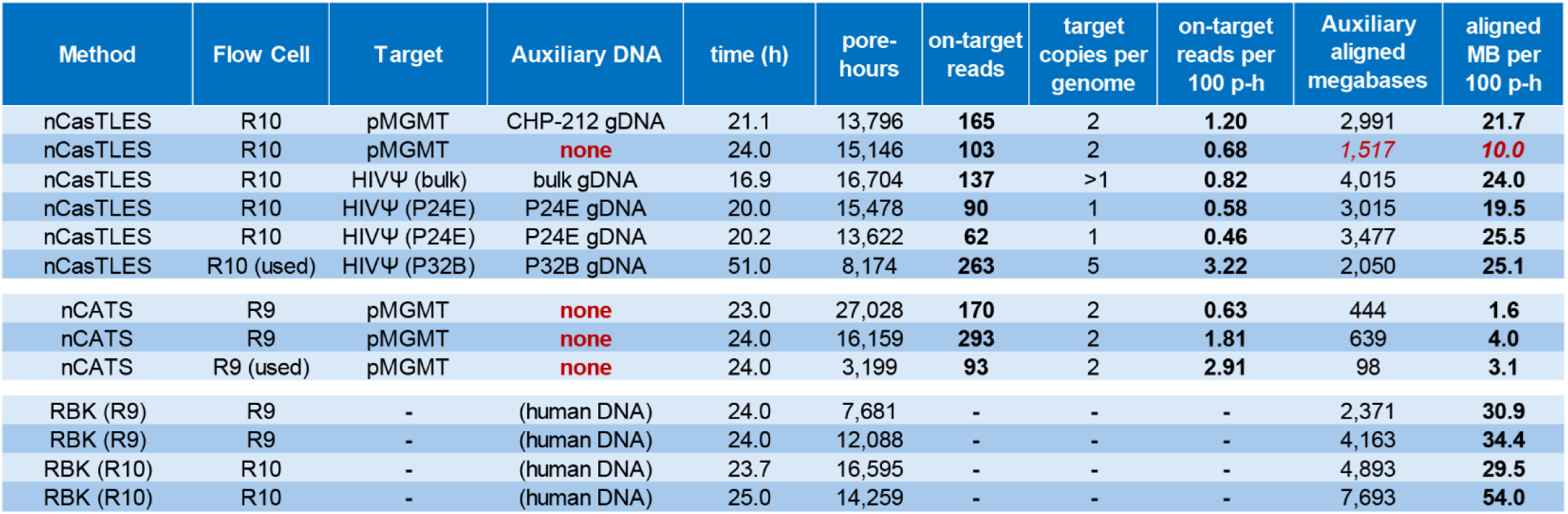
Sequencing run metrics. Table presents metrics from sequencing runs of nCasTLES on R10 flow cells, as well as data from previously published nCATS runs^14^ on R9 flow cells and non-enrichment strategy libraries on both flow cells. Pore-hours are calculated via the trapezoidal integration method from pore scan data during the run, e.g. a two-hour segment with a starting pore scan count of 200 pores and an end pore scan count of 100 pores would contain 150 pores * 2 hours = 300 pore-hours.

#### nCasTLES auxiliary DNA permits deeper off-target or parallel experimental sequencing

Efficient on-target read capture without loss of overall sequencing yield is not possible with current targeted nanopore sequencing methods like nCATS, but is possible with nCasTLES. Indeed, nCasTLES displayed similar on-target read counts as nCATS libraries previously run on R9 flow cells, while displaying greatly enhanced auxiliary read counts, approaching the throughput of a non-targeted nanopore sequencing run (**Table 1**). To demonstrate the utility of these reads, we analyzed the auxiliary CHP-212 gDNA that had been sequenced during a single run of the *MGMT* promoter (**Fig. 3a**). We had previously generated a map of the CHP-212 ecDNA using a standard rapid barcoding kit sequencing run and assembled the structure of the ecDNA using CoRAL (**Fig. 3b**). This assembly was identical to one generated using public CHP-212 sequencing data^11^. To assess the utility of auxiliary DNA sequencing, we next took the barcoded auxiliary reads from the MGMT sequencing runs and attempted to recreate the known structure of the ecDNA. Using these reads as input, CoRAL^17^ was able to successfully assemble the correct structure of the ecDNA (**Fig. 3c**). Thus, in the same sequencing run we were able to characterize the promoter of a disease-relevant gene in one set of cell lines and assemble the driver genomic alterations in a second cell line, demonstrating the utility of nCasTLES’ potential for multiplexing with other sequencing libraries.

**Figure 3.**
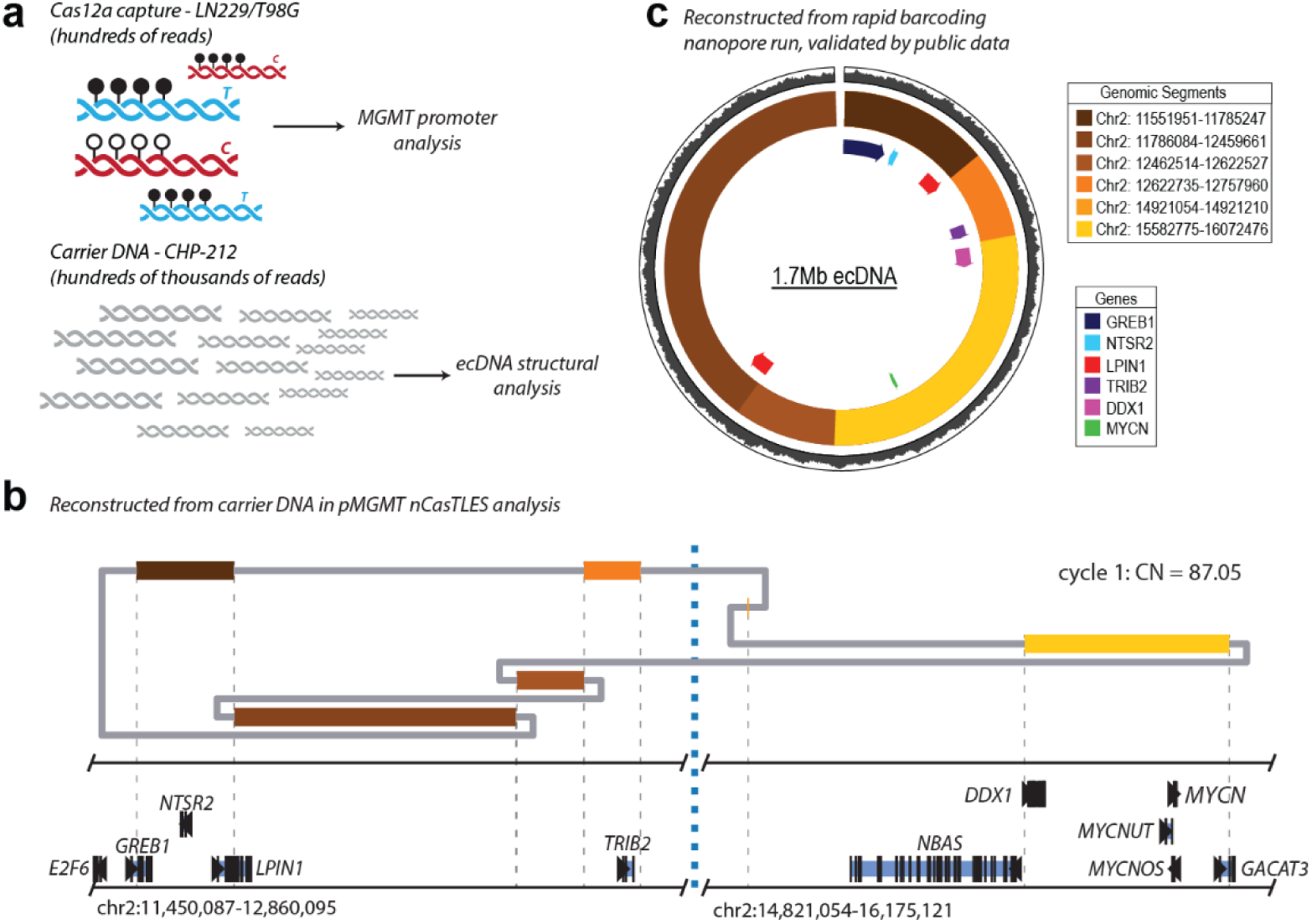
Auxiliary DNA reads permit characterization of extra-chromosomal DNA. (**a**) In one of the two sequencing experiments presented in Figure 2, nCasTLEs-enriched DNA was combined with CHP-212 auxiliary DNA for sequencing. While the on-target DNA permitted detailed interrogation of the *MGMT* promoter in GBM cell lines, the auxiliary CHP-212 DNA permits analysis of the ecDNA contained within those cells. (**b**) CHP-212 gDNA prepared via rapid barcoding kit and nanopore sequenced was used to construct a high-coverage map of the ecDNA element, which consists of six sections of chromosome 2. This constructed sequence matches one generated with publicly available sequencing data, and with previously published reports of the ecDNA sequence in this cell line. (**c**) Analysis of the CHP-212 auxiliary DNA sequenced during the MGMT promoter analysis resulted in construction of the same ecDNA structure.

### Single-cut capture of integrated lentiviral vector elements

We next sought to examine the single-cut potential of nCasTLES to profile DNA sites where only part of the sequence was known, such as the integration sites of a lentiviral transgene vector. Lentiviral integration sites into the genome display chromatin-state-specific biases^18,19^ but are otherwise effectively random and can serve as clone-specific barcodes. Traditional PCR-based integration site analysis (ISA) methods, like iPCR, can introduce amplification bias, remove DNA modifications, and require strategies like unique molecule identifiers/barcodes for full clonal interrogation^20^. nCATS-derived nanopore sequencing ISA methods, such as AFIS-seq, avoid amplification bias but suffer from the same pitfalls as nCATS; namely high input DNA requirements, rapid flow cell degradation and incompatibility with library multiplexing^21^. We reasoned that nCasTLES could be implemented for ISA and combined with whole-genome BrdU sequencing to explore the relationship between lentivirus integration sites and proliferation.

We infected U2OS 2-6-3 cells^22^ with a lentiviral rTTA expression vector that also contained a puromycin resistance cassette. U2OS 2-6-3 cells contain an integrated array consisting of CFP under the control of a tetracycline response element (TRE). rTTA expression will thus result in CFP expression under doxycycline, and infected cells will be resistant to puromycin regardless of dox (**Fig. 4a**).

**Figure 4.**
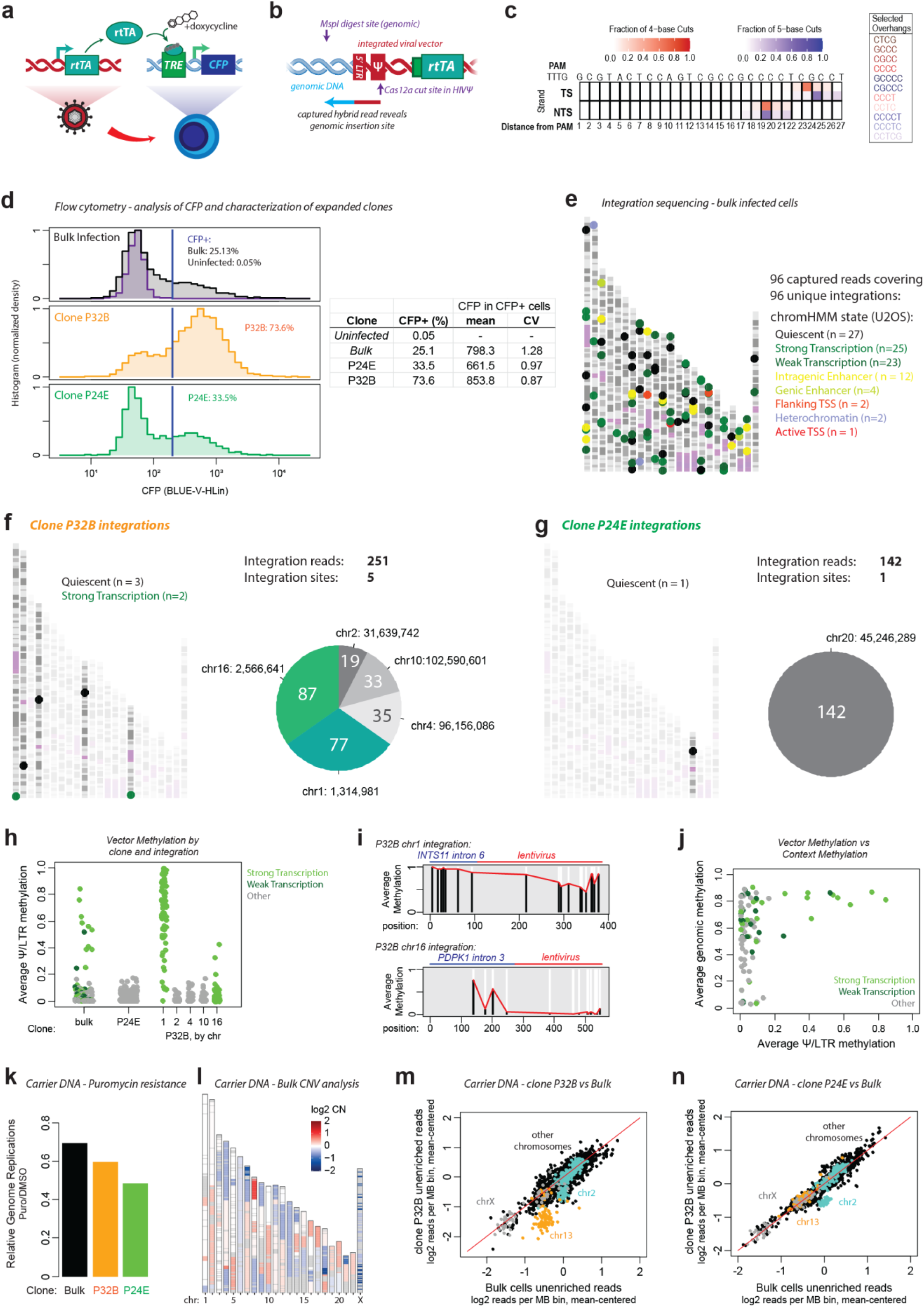
nCasTLES characterization of integrated lentiviral elements. (**a**) Schematic depicts experimental setup. U2OS 2-6-3 cells contain an integrated CFP locus downstream of a tet-responsive element array. Lentiviral infection with a reverse tetracycline-controlled transactivator (rtTA) causes doxycycline-mediated transcription of CFP. (**b**) Schematic depicts nCasTLES design. Cas12a guides targeting the HIVѰ sequence of the lentiviral vector were designed and validated. Genomic DNA from rtTA lentivirus-infected cells was captured with nCasTLES and cleaved with the MspI restriction enzyme, which would cleave at some point within the genomic DNA near the random lentiviral insertion, creating a sequencable fragment containing both the lentiviral 5’ LTR and some amount of surrounding genomic DNA. (**c**) Heatmap depicts results of HIVѰ Cas12a overhang characterization. Eleven predicted overhangs were tested as a pool and successfully enriched HIVѰ DNA (see Sup. Fig. 4). (**d**) Histograms depict flow cytometry characterization of CFP response in bulk infected 2-6-3 cells (black, top), as well as two selected clones, P32B (orange) and P24E (green). Table presents percentage of CFP positivity as well as mean and CV of CFP signal in CFP-positive cells from each population. (**e**) Plot depicts detected integration locations in bulk infected 2-6-3 cells, colored by chromHMM chromatin context. (**f**) Plot depicts five integration sites detected in clone P32B, as well as chromatin contexts. Pie chart depicts distribution of read counts covering each detected site. (**g**) Plot depicts the single integration site detected in clone P24E, as well as its chromatin context. Pie chart depicts distribution of read counts covering the detected site. (**h**) Plot depicts methylation of CpGs within the lentiviral element for each read, by clone and integration site. Points are colored based on whether they are in a transcription chromHMM state or not. (**i**) Trace depicts average methylation at each CpGs within reads from the chr1 (top) and chr16 (bottom) integration sites in clone P32B. Each vertical line represents a single CpG, with black and white representing the methylated and unmethylated proportions, respectively. The red line depicts the average methylation across the read. The horizontal red and blue lines above each trace depict the lentiviral and genomic/intronic sections of each read. (**j**) Plot depicts lentiviral and context methylation for each bulk read, colored by whether the integration site was within a transcription chromHMM region or not. (**k**) Bar plot depicts relative genome replications detected for each clone under 20ug/mL puromycin compared to DMSO control, detected by BrdU positivity. Relative genome replications are calculated from BrdU positivity as: BrdU+ / (1-BrdU+). (**l**) Plot depicts relative copy number status from auxiliary bulk 2-6-3 reads, with blue showing regions of loss and red showing amplified regions. Intensity of color corresponds to degree of loss/amplification. (**m,n**) Plot depicts read density from auxiliary sequencing in 1MB bins across the entire genome, from P32B vs bulk sequencing (m) or P24E vs bulk sequencing (n). Points are log2 scale and mean-centered. Bins from chr13 are depicted in orange, bins from chr2 are depicted in light blue, bins from chrX are depicted in gray, and other bins are depicted in black. Red lines depict the diagonal (y=x), that is, similar bin coverage in each condition.

We designed a Cas12a guide targeted to the HIV-ψ sequence (**Fig. 4b**) using CRISPOR^12^. We confirmed efficient digestion by qPCR, then designed and validated eleven capture overhang sequences as described above (**Fig. 4c, Sup. Fig. 4**). We infected U2OS 2-6-3 cells at a target MOI of 1, selected in 10ug/ml puromycin for 48 hours, and then analyzed cells via flow cytometry. Bulk infected cells showed a strong CFP response to 500 ng/ml doxycycline (**Fig. 4d**). We selected several CFP^+^ clones by flow sorting for expansion and characterization, including clones designated P24E and P32B. When expanded, both clones displayed robust CFP signal following doxycycline induction. Notably, P32B displayed a higher positivity rate and signal intensity from positive cells than P24E, and both clones displayed a lower coefficient of variance in the fluorescence intensity of CFP^+^ cells than the parental bulk population (**Fig. 4d**). We hypothesized that higher vector copy number and integration into methylated introns of highly transcribed genes improve the expression of delivered transgenes.

We also sought to utilize the auxiliary DNA to more deeply characterize the selected clones in two ways. First, we sought to determine if the location of lentivirus integrations and/or the vector copy number could affect puromycin resistance, given that the puromycin resistance cassette was part of the lentiviral vector. To do so, we harnessed BrdU incorporation to track genome replications as a surrogate for proliferation. Second, we sought to utilize the auxiliary DNA as whole-genome sequencing data to track any chromosomal copy number alterations that occurring during clonal selection and outgrowth. To accomplish both aims, bulk infected U2OS 2-6-3 cells and cells from each clone were grown in either 10uM BrdU alone or 10uM BrdU and 20ug/mL puromycin for 24 hours, which is a dose higher than used for initial selection. Genomic DNA from each culture was cut with the ψ-specific guide and the eleven capture oligos were ligated to this cut DNA. DNA was then digested with MspI to fragment the genomic DNA, and captured fragments were isolated via streptavidin beads. Thus, each sequenced library contained nCasTLES-enriched lentiviral integrations as well as auxiliary DNA containing BrdU, allowing us to simultaneously profile (1) the lentiviral integrations, (2) genomic replications as a readout of the functional response of the lentiviral puromycin resistance, and (3) total genomic coverage as a readout of chromosomal copy number alterations.

For the bulk infected cells, nCasTLES captured 137 total reads that mapped to the 5’LTR (**Table 1**), 100 of which also mapped to hg38 at 96 unique sites (**Fig. 4e**). Closer examination of the four sites with two reads revealed that these reads were duplex pairs^23^, indicating that each of these sites had two reads of opposite strand DNA sequenced sequentially through the same pore and were thus opposite strands of the same captured molecule (**Sup. Fig. 5**). Thus, with 96 unique molecules and no duplicate captures, the 95% confidence interval of maximum clonal representation is no more than 3.2% by the rule of three^24^, and our bulk population represents a highly heterogenous lentiviral infections. For P32B, we captured 251 reads, covering 5 distinct integration sites with between 19 and 87 reads (**Fig. 4f**). Notably, the read count was inversely correlated with predicted molecular fragment size based on the distance between the integration site and the nearest genomic MspI digestion site; the variance in read counts thus likely represents capture efficiency rather than differences in representation (**Sup. Fig. 6**). Of these sites, three were in areas considered “quiescent” by chromHMM analysis^25,26^, while two were in the introns of highly expressed genes. For P24E, we captured 142 lentiviral integration molecules, all of which mapped to the same site on chromosome 20, which was categorized as “quiescent” chromHMM (**Fig. 4g**). These data demonstrate the utility of nCasTLES as a viable strategy for ISA.

#### Epigenetic analysis of lentiviral integrations and surrounding genomic DNA

One additional advantage of nCasTLES is that DNA modification information is preserved, which enabled us to explore the methylation profile of the lentiviral 5’ LTR and ψ sequences as well as that of surrounding genomic DNA. Modern lentiviral vectors are engineered with elements to prevent transgene methylation and with self-inactivating mutations in the 3’ LTR that remove the functional promoter/enhancer U3 elements, which are copied to the 5’ LTR during integration, so methylation of the captured lentiviral sequence is unlikely to affect transgene expression or lentiviral activity^27,28^. However, we reasoned that methylation examination could potentially offer insight into the dynamics of methylation regulation. We used remora^29^ to call methylation at the 12 CpGs within the lentiviral regions (the 5’ LTR and part of the HIVψ) in our captured reads. Little to no methylation was detected in any of the P24E captured reads, in the majority of bulk integration sites, nor in four of the five P32B integration sites (**Fig. 4h**). We detected strong methylation in the chr1 P32B integration site, and in around 10% of the bulk integration sites. Notably, while not all integrations within strongly transcribed regions became methylated, all integration sites that became methylated were located in transcribed regions of the genome. We explored this further with the two P32B transcribed integration sites by analyzing the methylation within the genomic and lentiviral sections of the reads (**Fig. 4i**). The P32B chr1 integration site, which became methylated, was inserted into the sixth intron of the gene *INTS11* and located near a cluster of methylated CpGs. This methylation appears to have successfully spread into the lentiviral integration. In contrast, at the P32B chr16 integration site within the third intron of the gene *PDPK1*, there are fewer nearby CpG sites and these sites display less methylation in our captured reads. Consistently, in analysis of the bulk reads, all reads that displayed methylation in the lentiviral CpGs also displayed strong methylation in nearby captured genomic CpGs, but again, not all genomically-methylated reads gained lentiviral methylation (**Fig. 4j**). These data suggest that nearby genomic methylation is necessary but not sufficient for lentiviral methylation, at least at this analyzed timescale.

#### Utilization of U2OS 2-6-3 auxiliary DNA for genomic and functional characterization of infected cells

We postulate that the increased number of integration sites and/or having integration sites in highly transcribed regions may result in higher transgene expression, conferring a higher doxycycline-induced CFP response to clone P32B in comparison to clone P24E (as in Fig. 4e). We next sought to explore the effects of these different lentivector integrations on resistance to puromycin, by analyzing the auxiliary DNA BrdU incorporation to track genome replications, as a surrogate for proliferation rate. BrdU incorporation was called using a custom Remora model validated by barcoded split-pool synthesized BrdU training libraries^29,14^. We hypothesized that the higher vector copy number and vector integration into highly transcribed regions would confer higher transgene expression and puromycin resistance in clone P32B cells compared to clone P24E cells. Therefore, we reasoned that clone P32B cells should exhibit faster growth and, as a result, more frequent BrdU-positive molecules than P24E cells under puromycin treatment. Indeed, we found that clone P32B cells had a higher fraction of BrdU positive reads under puromycin treatment than P24E cells (**Fig. 4k**), a finding which was validated via cell viability assay (**Sup. Fig. 7b**).

We sought to use the auxiliary DNA to examine any large-scale chromosomal copy-number changes that developed during selection and outgrowth of the clones. Using QDNAseq^30^, we calculated relative chromosomal copy numbers present in the bulk population (**Fig. 4l**). As our integration analysis demonstrated that no single clone could represent more than 3% of the bulk population, we reasoned this was likely an accurate background state to compare clonal chromosomal copy number changes to. Copy number variations (CNVs) were evaluated in the auxiliary DNA from bulk versus each clone using QDNAseq in one megabase bins. In clone P32B, we consistently saw that bins in chromosome 13 fell below the diagonal and had less relative reads in P32B than in the bulk population, suggesting a loss of a chr13 copy in that clone. We also observed several bins at the Q end of chr2 fall slightly below the diagonal (**Fig. 4m**). In comparison, chromosome X displayed low counts in both bulk and P32B, but the relative counts were similar and the bins stayed close to the diagonal. We saw no signature of chr13 gain or loss in clone P24E and most bins were close to the diagonal, with the exception of the same bins in at the Q end of chr2 (**Fig. 4n**).

These data demonstrate the unique ability of nCasTLES to profile targets of interest in which only part of the sequence of the target is known, and to profile genetic and epigenetic information from those reads. Further, nCasTLES allows for simultaneous targeted sequencing and profiling of genome-wide DNA modifications, such as BrdU incorporation, and was able to demonstrate a presumed loss of a chr13 copy during selection and outgrowth of one of the interrogated clones, demonstrating the versatility and efficiency of the method and the power of simultaneous targeted DNA and auxiliary DNA sequencing.

## Discussion

Here, we present nCasTLES, an updated Cas12a-based method for the sequence-specific isolation of native DNA for applications such as nanopore sequencing. By targeted ligation to characterized overhangs generated by Cas12 cleavage, nCasTLES allows for enrichment of native DNA rather than preferential activation, and is thus an adaptable methodological framework for targeted nanopore sequencing without loss of flow cell throughput.

One of the strengths of our method is the ability to spike nCasTLES-enriched libraries into other libraries to increase flow cell throughput. nCATS results in a high number of on-target reads per sequencing run, however one significant drawback is the dramatically reduced total throughput of the flow cell. This does serve to make the on-target read fraction higher; the numerator (on-target reads) is increased while the denominator (off-target reads) is decreased, however this comes at the cost of lower overall yield of potentially useful sequencing data. For instance, here we present uses such as characterization of BrdU incorporation, assessment of chromosomal copy number variations during clonal outgrowth, and assembly of the structure of an ecDNA element. Thus, the increased off-target read capacity from nCasTLES allows for more efficient data generation and can enable unique experimental designs.

Our method is well-suited for the characterization of disease-relevant genomic methylation, such as the promoter of *MGMT. MGMT* methylation status in glioblastoma is predictive of clinical response to the frontline chemotherapeutic temozolomide. However, PCR amplification and bisulfite sequencing of this promoter are challenging due to high GC content and limited abundance of clinical material. Direct profiling of CpG methylation at this locus with targeted third-generation sequencing would overcome these hurdles. We demonstrate that nCasTLES is well-suited for isolation and characterization of the MGMT promoter and show that it can characterize subtle differences in the methylation profile of the promoter and surrounding intragenic/intronic CpGs through mixing of cell lines with distinct methylation dynamics. Unsupervised clustering of the data generated through nCasTLES was able to correctly distinguish both the methylation state of the promoter and the cell-line of origin of the read. Additionally, the high-depth coverage and full-locus single-molecule profiling allowed for the identification of potentially interesting subpopulations, such as the unmethylated promoters that appeared to be in a transitory state with encroaching methylation; these rarer molecules would not have been identifiable with shorter-read sequencing approaches.

An additional advantage of nCasTLES is its ability to function with a single cut site to isolate unknown DNA. Here, we isolated integrated lentiviral vector elements and captured the surrounding genomic DNA. Traditionally, isolation of the integration site of a lentivirus would require PCR^20^, which could introduce amplification biases and would prevent characterization of lentiviral or contextual genomic methylation. Our approach is amplification-free and allows for the accurate estimation of clonality of a population, as well as an exploration of methylation dynamics. Additionally, since nCasTLES can be spiked into another library for sequencing, this estimation of clonality could be performed on top of another characterization, such as the BrdU incorporation following selection or assessment of clonal-selection associated chromosomal alterations we performed here.

Notably, there are some significant limitations to nCasTLES, as presented in this manuscript. First, the bead-based capture does seem to impose a limit on the upper length of molecules that can be efficiently isolated. nCasTLES with bead enrichment may not be suitable for applications requiring very long (>25-50kb) read lengths. We anticipate that modifications to skip the bead step may prove useful here; for instance, utilizing two Cas12 cut sites on either side of a large fragment, ligation of hairpin adaptors to each end, and then exonuclease digestion of unprotected DNA. Post-exonuclease cleavage of one of these hairpins, either through the addition of a sequence-specific endonuclease motif or through cleavage of a non-standard base incorporated into the hairpin (e.g. deoxyuracil cleaved by uracil-DNA glycosylase/USER Enzyme^31^ or deoxyinosine cleaved by Endonuclease V^32^) could reveal a ligation site for library preparation. Additionally, as presented we utilize a sequence-specific restriction endonuclease to free enriched DNA from the beads, which imposes an upper limit on fragment size based on recognition site frequency, but a similar non-standard base cleavage approach within the biotinylated tether oligo could also be employed here. As presented, nCasTLES also requires characterization of the site-specific overhangs left by Cas12a cleavage. The main function of this is to prevent adapter self-ligation; here we utilize between two and eleven overhangs, but a larger pool of overhangs would not present problems if adapter self-ligating sequences were omitted. Combined with the development of better predictive methods regarding the Cas12a cleavage preference, this step could become unnecessary.

In summary, nCasTLES presents a simple framework for enhanced nanopore-based target isolation and sequencing, with the added benefit of being adaptable as needed to enhance specific experimental objectives.

## Supporting information

Supplemental Note 1 nCasTLES protocol

## Acknowledgements

We thank D. Davis, S. David, K. Droppa, E. Watson, and S. Wolfe for thoughtful discussion. We thank J. Benanti, S. Cantor, and T. Fazzio for thoughtful discussion and helpful comments on the manuscript. We thank L. Robbins and the UMass Chan HPC administrators for computational support. We thank D. Spector (CSHL) for the U2OS 2-6-3 cell line.

## Author Contributions

Conceptualization: MV, KK, WAF

Methodology: MV, KK, BAP, WAF

Investigation: MV, KK, BAP

Visualization: MV, WAF

Supervision: WAF

Writing: MV, WAF

## Data and materials availability

All processed data are presented in the main text or the supplementary materials and data. Raw sequencing reads (fastq files) are available in SRA under accession PRJNA1518188.

## Supplementary Materials

Figures S1 through S7.

Supplementary Note 1: detailed nCasTLES protocol.

## Materials and Methods

### Design of Cas12a guides

We obtained hg38 sequences for the *MGMT* locus and for the *MYCN* locus from the UCSC genome browser as fasta files. The HIV-ψ sequence was obtained from the Addgene plasmid map of pLentiCRISPR_SDHA (Addgene, Plasmid #177980). Cas12a guides for all three targets were designed with CRISPOR^12^ and synthesized by IDT.

MYCN_guide: rUrArArUrUrUrCrUrArCrUrArArGrUrGrUrArGrArUrArArUrUrUrGrArArArCrUrGrGrArArA rCrArUrCrCrArG,

MGMT_guide: rUrArArUrUrUrCrUrArCrUrArArGrUrGrUrArGrArUrGrUrGrUrCrCrUrArArGrArArArGrGrG rUrGrGrGrArArG

HIV-ψ_guide: rUrArArUrUrUrCrUrArCrUrArArGrUrGrUrArGrArUrGrCrGrUrArCrUrCrArCrCrAr GrUrCrGrCrCrGrCrCrCrC.

### Cell Culture

CHP-212 (ATCC, CRL-2273) cells were cultured in DMEM/F12 with 10% FBS and pen/strep. T98G (ATCC, CRL-1690) and LN229 (ATCC, CRL-2611) cells were cultured in DMEM with 10% FBS and pen/strep. U2OS 2-6-3 cells were a kind gift from David Spector and were cultured in DMEM with 10% FBS and pen/strep.

### CHP-212 ecDNA Reconstruction

Genomic DNA from CHP-212 cells was extracted via Monarch Spin gDNA Extraction Kit (NEB), prepared for sequencing via Rapid Barcoding Kit 24 V14 (Oxford Nanopore Technologies), and sequenced on an R10 MinION Flow Cell (DNA) for 72 hours.

Basecalling was performed with Dorado v2.0.0^34^ with model dna_r10.4.1_e8.2_400bps_fast@v5.0.0. Reads were aligned to hg38 using the minimap2 suite.^35^ Coverage tracks were generated with Deeptools bamCoverage, with normalization by 1x depth (reads per genome coverage, RPGC).^37^ ecDNA reconstruction was performed with CoRAL^39^ and data were plotted using the R package circlize^40^.

### Validating Performance of MYCN Cas12a guide

A 1603bp PCR product covering the MYCN guide target site was generated from CHP-212 gDNA using Q5 High-Fidelity DNA Polymerase (NEB) and PCR primers for the MYCN promoter (MYCN_prom_full_F: TCCCCAGAAGAATAGCATGTCG, MYCN_prom_full_R: GCTGCAAAAGGATTAGGGCG), purified via gel extraction with Monarch Spin DNA Gel Extraction Kit (NEB), and was digested with Cas12a ribonucleoproteins (RNP) loaded with *MYCN* gRNA. Digestion efficiency was assessed with agarose gel electrophoresis **(Sup. Fig. 1a).**

On-target cleavage was validated in CHP-212 genomic DNA; 50ng, 100ng or 200ng of gDNA was digested with *MYCN*-targeting Cas12a RNP or RNP-free negative controls. qPCR was performed with 2x AzuraView GreenFast qPCR Blue Mix LR (Azura Genomics) and primers for GREB1 (GREB1_F1: GGCCTGTGAGGCTGGTTATT, GREB1_R1: AGGCATCTGACTAGCTCCCA), the MYCN Promoter (MYCN_promoter_F2: CCCTGCTATTTTGCACCTTCG, MYCN_promoter_R2: AGGGGAGACCGATGCTTCTA) and the MYCN Cas12a cut site (MYCN_Cas12_cut_1F: ACCCTCGTAGCTCGCACTTA, MYCN_Cas12_cut_1R: GGCTAGTCCGAAGGTGCAAA) **(Sup. Fig. 1b).**

### Profiling Cleavage Sites Generated by MYCN-targeting Cas12a RNP

CHP-212 genomic DNA was digested with MYCN-targeting Cas12a RNP and ligated to annealed DNA adapters containing variable 4N- or 5N-5’ overhangs in separate reactions (Amp_seq_F: TGGAAAGGATTCATTCCCACGGCTTGGGATGAATGGCAACTGTCGAGGACCTATCCACTAGACGAGTATTGCTGCAGTCTGCA), (Reverse adapters: Amp_seq_4N_R: /5Phos/NNNNTGCAGACTGCAGCAATACTCGTCTAGTGGATAGGTCCTCGACAGTTGCCATTCATCCCAAGCCGTGGGAATGAATCCTTTCCA, Amp_seq_5N_R: /5Phos/NNNNNTGCAGACTGCAGCAATACTCGTCTAGTGGATAGGTCCTCGACAGTTGCCATTCATCCCAAGCCGTGGGAATGAATCCTTTCCA). Excess adapters were removed with 0.4X AMPure XP bead size selection. Regions spanning the Cas12a cut site were amplified by PCR using a primer for the PCR adapter (bc7_qPCR_R: GGAAAGGATTCATTCCCACGG) and a locus-specific primer (MYCN_promoter_R: AGCGCGTCCAGACAGATGAC) using Q5 High-Fidelity DNA Polymerase (NEB). Amplified products were purified using 0.5X AMPure XP bead cleanup and submitted for PCR-EZ amplicon sequencing (Genewiz). To identify the sequences of the Cas12a overhangs, CUTADAPT^1^ was first deployed to identify and remove the 4N- and 5N-adapters from the fastq files. The sequences of the 4 or 5 bases remaining after adapter removal were extracted with awk. Sequences not found in the target region were filtered out. These sequences were used to design 4 reverse oligos with 5’ overhangs capable of ligation to the target side of the MYCN Cas12a cut site (MYCN Reverse adapters: MYCN_1R: /5Phos/ TCTGGCCAAGACCTGCAGGGGATCCTGATCACCATGGGAGCTCCCTAGGCCAC, MYCN_2R: /5Phos/ CTGGCCAAGACCTGCAGGGGATCCTGATCACCATGGGAGCTCCCTAGGCCAC, MYCN_3R: /5Phos/ TGGATCCAAGACCTGCAGGGGATCCTGATCACCATGGGAGCTCCCTAGGCCAC, MYCN_4R: /5Phos/GGATCCAAGACCTGCAGGGGATCCTGATCACCATGGGAGCTCCCTAGGCCAC).

### Validating MYCN Promoter Enrichment with biotinylated DNA adapters

Biotinylated DNA adapters, consisting of a forward biotinylated oligo (SbfI_biotin_TEG_F: /5BiotinTEG/TGGCAACTGTCGGAGGACCTATCCACTAGACGAGGTAATTCCTGCAGGT CTTGG) annealed to each of the MYCN-specific reverse adapters, described above, were each separately ligated to Cas12a-digested *MYCN* PCR product. Excess adapter was removed with 0.6X Ampure XP bead size selection, remaining DNA was then bound to Pierce Streptavidin beads (Thermo Scientific). Unbound DNA was removed with washes in 2X and 1X SSC Buffer (Thermo Scientific), followed by washes with nuclease-free water. Target DNA was released from the beads with SbfI-HF (NEB) digest, then purified with the Monarch Spin PCR & DNA Cleanup Kit (NEB). The target DNA was quantified with the Qubit 1X dsDNA High Sensitivity (HS) Assay Kit (Invitrogen) (**Sup. Fig. 1C**, **Fig. 1D)**. Captured target size was assessed with agarose gel electrophoresis (**Sup. Fig. 1D).** Adapters MYCN_1R, MYCN_2R were selected for further testing.

High molecular weight gDNA was extracted from CHP-212 cells via Monarch HMW DNA Extraction Kit (NEB), then digested with MYCN-targeting Cas12a RNP. Cas12a digests were purified with Monarch Spin PCR & DNA Cleanup Kit (NEB). Biotinylated DNA adapters with MYCN_R1 and MYCN_R2 were ligated together to the Cas12a digest, followed by purification with 0.6X Ampure XP bead selection. 5uL of the ligation reaction was used as the no pulldown control. Target DNA was enriched from the remainder of the ligation reaction with Pierce Streptavidin bead pulldown and SbfI-HF release as previously described. qPCR was performed with 2x AzuraView GreenFast qPCR Blue Mix LR (Azura Genomics) and primers for GAPDH (non-target DNA) (GAPDH-early-genic-qPCR_F: AAGGAGAGCTCAAGGTCAG, GAPDH-early-genic-qPCR_R: AGTAGGGACCTCCTGTTTCT), the MYCN Promoter (Target DNA) (MYCN_promoter_F2: CCCTGCTATTTTGCACCTTCG, MYCN_promoter_R2: AGGGGAGACCGATGCTTCTA), and a site upstream of the MYCN Cas12a cut site on the ecDNA (UP_MYCN_F1: AGTTCCAGGAGCCAAAGAGC, UP_MYCN_R1: ATCGAACTCTGCACTGGTGG).

### Targeted Sequencing of ecDNA MYCN locus from CHP-212

nCasTLES was performed to generate the MYCN ecDNA enrichment library as follows: CHP-212 gDNA was digested with MYCN-targeting Cas12a RNP and purified via Monarch Spin PCR & DNA Cleanup Kit (NEB). Biotinylated DNA adapters with MYCN_R1 and MYCN_R2 were ligated to the Cas12a digest, followed by purification with 0.6X Ampure XP bead size selection. Target DNA was enriched from the ligation reaction with Pierce Streptavidin bead pulldown and SbfI-HF release as described above. A non-enrichment control library was generated by restriction digest of the CHP-212 gDNA with SbfI-HF, followed by 0.4X AMPure XP bead cleanup. 16.7ng of the MYCN ecDNA enrichment library and 46.2ng of the non-enrichment control library were combined and prepared for sequencing via Ligation Sequencing Kit V14 (ONT). The library was sequenced on a MinION R10 Flow Cell (ONT) for 25 hours and 47 minutes. In a separate run, 62.9ng of the non-enrichment control library was prepared for sequencing via Ligation Sequencing Kit, then sequenced on an R10 Flow Cell for 72 hours.

Basecalling was performed with Dorado v2.0.0^34^ with model dna_r10.4.1_e8.2_400bps_fast@v5.0.0. Reads were aligned to hg38 using the minimap2 suite^35^. Bedtools intersect was used to identify reads aligned to the MYCN target fragment^36^. Coverage tracks were generated with Deeptools bamCoverage, with normalization by 1x depth (reads per genome coverage, RPGC)^37^. Coverage plots were generated using the R package plotgardener^38^. ecDNA reconstruction was performed with CoRAL^39^.

### Validating Performance of MGMT Cas12a guide

A 1627bp PCR product covering the MGMT guide target site was generated from LN-229 gDNA using Q5 High-Fidelity DNA Polymerase (NEB) and PCR primers for the MGMT promoter (MGMT_G4_Amp_F: GAAGTAGACCACCAGTACGGA, MGMT_Full_2R: TCAAAACAAGAGTGGGGCGA). The PCR product was purified via Monarch Spin PCR & DNA Cleanup Kit (NEB). The PCR product was digested with MGMT-targeting Cas12a RNP. Digest was assessed with agarose gel electrophoresis **(Sup. Figure 2A).**

### Profiling Cleavage Sites Generated by MGMT-targeting Cas12a RNP

LN229 gDNA was digested with *MGMT*-targeting Cas12a RNP, ligated to 4N- and 5N-overhang adapters, PCR amplified as described above except with an *MGMT* locus-specific primer replacing the *MYCN* primer (MGMT_G4_Amp_F: GAAGTAGACCACCAGTACGGA), and sequenced. Based on the sequencing results, candidate reverse oligos with 5’ overhangs capable of ligation to the target side of the MGMT Cas12a cut site were designed: (MGMT Reverse adapters: MGMT_G4_R1: /5Phos/CTTCCCAAGACCTGCAGGGGATCCTGATCACCATGGGAGCTCCCTAGGCCAC, MGMT_G4_R2: /5Phos/TTCCCCAAGACCTGCAGGGGATCCTGATCACCATGGGAGCTCCCTAGGCCAC, MGMT_G4_R3: /5Phos/TCCCCCAAGACCTGCAGGGGATCCTGATCACCATGGGAGCTCCCTAGGCCAC, MGMT_G4_R4: /5Phos/CTTCCCCAAGACCTGCAGGGGATCCTGATCACCATGGGAGCTCCCTAGGCCAC, MGMT_G4_R5: /5Phos/TCTTCCCAAGACCTGCAGGGGATCCTGATCACCATGGGAGCTCCCTAGGCCAC, MGMT_G4_R6: /5Phos/TTCCCCCAAGACCTGCAGGGGATCCTGATCACCATGGGAGCTCCCTAGGCCAC).

### Validating MGMT Promoter Enrichment with biotinylated DNA adapters

The 700bp target fragment was gel purified from Cas12a RNP-digested MGMT PCR product. Biotinylated DNA adapters were made by annealing the forward biotinylated oligo, as above, to each of the *MGMT*-specific reverse adapters in separate reactions, and purified as above, except target DNA was released from the beads with NcoI-HF (NEB) digest (**Sup. Fig. 2b,c**). Adapters MGMT_G4_R1, MGMT_G4_R2, MGMT_G4_R3, MGMT_G4_R4 were selected for further testing.

To create no-pulldown controls, LN229 HMW gDNA or T98G HMW gDNA was digested with NcoI-HF and PflMI (NEB), followed by purification. The pulldown library was made as follows: LN229 HMW gDNA or T98G HMW gDNA was digested with MGMT-targeting Cas12a RNP, followed by heat inactivation at 65C. Biotinylated DNA adapters with MGMT_G4_R1, MGMT_G4_R2, MGMT_G4_R3 and MGMT_G4_R4 were ligated to the Cas12a digest directly in the Cas12a reaction mix with T4 DNA Ligase (NEB, M0202L), followed by heat inactivation at 65C. The ligation was digested with PflMI. Streptavidin pulldown was performed with the Dynabeads kilobaseBINDER Kit (Invitrogen). Target DNA was released from the beads with NcoI-HF digest, then purified with the Monarch Spin PCR & DNA Cleanup Kit (NEB). qPCR was performed with primers for an area downstream of MYCN (non-target DNA) (DOWN_MYCN_F1: CTCTGTCCCTTAAGGAGCAACC, DOWN_MYCN_R1: TGTACAGGTGCAAAAAGGGCT), the cell line SNP within the MGMT Promoter (target DNA) (MGMT_SNP_F: TGGGTAGCACCTTGTTCAGC, MGMT_SNP_R: TGAGACACCTGGGGAGGAAA), and a second area in the MGMT promoter (target DNA) (MGMT_G5_cut_F: CAGGACCGGGATTCTCACTAAG, MGMT_G5_cut_R: CTTAGTTTGCCAAATGGCCCG) **(Sup. Fig 2d**).

### Targeted Sequencing of MGMT Promoter from LN229 and T98G cells

nCasTLES was performed to generate the MGMT enrichment libraries as follows: LN229 gDNA or T98G gDNA digested with MGMT-targeting Cas12a RNP, followed by heat inactivation. Biotinylated DNA adapters with MGMT_G4_R1, MGMT_G4_R2, MGMT_G4_R3 and MGMT_G4_R4 were ligated to the Cas12a digest directly in the Cas12a reaction mix with T4 DNA Ligase (NEB), followed by heat inactivation. The ligation was digested with PflMI. Streptavidin pulldown was performed with the Dynabeads kilobaseBINDER Kit (Invitrogen). Target DNA was released from the beads with NcoI-HF digest, then purified. For the no-auxiliary DNA library, 6.44ng of the T98G MGMT nCasTLES library and 5.83ng of the LN229 MGMT nCasTLES library were combined and prepared for sequencing following the using the Ligation Sequencing Kit V14 (ONT), then sequenced on a MinION R10 Flow Cell (ONT) without auxiliary DNA for 24 hours. For the auxiliary DNA library, 6.302ng of the T98G MGMT nCasTLES library and 5.704ng of the LN229 MGMT nCasTLES library were combined and purified, then 1.28ng of the combined T98G/LN229 MGMT nCasTLES library and 17.6ng of the auxiliary DNA were combined and sequenced on an R10 flow cell for 21 hours and 8 minutes.

Sequenced data were processed and visualized as above. 5mCG calling was performed with Remora software from ONT using a custom model.

### Validating Performance of HIV-ψ Cas12a guide

pLentiCRISPRv1_SDHA (Addgene, Plasmid #177980) was digested with HIV-ψ-targeting Cas12a RNP. Digest was assessed with agarose gel electrophoresis (**Sup. Fig. 3a**).

### Profiling Cleavage Sites Generated by HIV-ψ-targeting Cas12a RNP

DNA adapters containing variable 4N- or 5N-5’ overhangs were annealed and each ligated in separate reactions to pLentiCRISPRv1_SDHA digested with HIV-ψ-targeting Cas12a RNP. Excess adapters were removed with 0.4X AMPure XP bead size selection. Regions spanning the

Cas12a cut site at the HIV-ψ locus were amplified by PCR using using a primer for the PCR adapter (bc7_qPCR_R: GGAAAGGATTCATTCCCACGG) and a locus-specific primer (M13/pUC Reverse: AGCGGATAACAATTTCACACAGG) using Q5 (NEB). Amplified products were purified using 0.4X AMPure XP bead size selection and sequenced. Candidate reverse oligos with 5’ overhangs capable of ligation to the target side of the HIV-ψ Cas12a cut site were designed as previously described (HIV-PSI-R1:

/5Phos/CGAGCCAAGACCTGCAGGGGATCCTGATCACCATGGGAGCTCCCTAGGCCAC, HIV-PSI-R2: /5Phos/
GGGCCCAAGACCTGCAGGGGATCCTGATCACCATGGGAGCTCCCTAGGCCAC, HIV-PSI-R3: /5Phos/
GGCGCCAAGACCTGCAGGGGATCCTGATCACCATGGGAGCTCCCTAGGCCAC, HIV-PSI-R4: /5Phos/
GGGGCCAAGACCTGCAGGGGATCCTGATCACCATGGGAGCTCCCTAGGCCAC, HIV-PSI-R5: /5Phos/
GGGGCCCAAGACCTGCAGGGGATCCTGATCACCATGGGAGCTCCCTAGGCCAC, HIV-PSI-R6: /5Phos/
GGGCGCCAAGACCTGCAGGGGATCCTGATCACCATGGGAGCTCCCTAGGCCAC, HIV-PSI-R7:
/5Phos/AGGGCCAAGACCTGCAGGGGATCCTGATCACCATGGGAGCTCCCTAGGCCAC, HIV-PSI-R8:
/5Phos/GAGGCCAAGACCTGCAGGGGATCCTGATCACCATGGGAGCTCCCTAGGCCAC, HIV-PSI-R9:
/5Phos/AGGGGCCAAGACCTGCAGGGGATCCTGATCACCATGGGAGCTCCCTAGGCCAC, HIV-PSI-R10:
/5Phos/GAGGGCCAAGACCTGCAGGGGATCCTGATCACCATGGGAGCTCCCTAGGCCAC, HIV-PSI-R11:
/5Phos/CGAGGCCAAGACCTGCAGGGGATCCTGATCACCATGGGAGCTCCCTAGGCCAC).

### Lentiviral Infection and Fluorescent Activated Single Cell Sorting

U2OS-2-6-3 cells (a gift from David Spector) were cultured in DMEM with 10% Tet-free FBS (Takara and Gibco) and pen/strep. 30 to 60 μL of lentiviral particle containing rtTA-M2 (GenTarget) with puromycin resistance gene (1 × 108 IFU/ml) were added to U2OS 2-6-3 cells in a 24-well plate. The next day, each well was pooled and moved to 6 wells, where 2ug/mL of puromycin was added to select for infected cells (Bulk Unsorted). To isolate single cell clones of U2OS 2-6-3 with different degrees of doxycycline-inducible CFP expression, 500ng/mL of doxycycline (MedChemExpress) was added to the media for 24 hours to induce CFP. Trypsinized cells were washed in PBS once, resuspended in PBS with 10% Tet-free FBS, and filtered before FACS. Cells were gated for FSC/SSC for intact cells, FSC-A/FSC-H for singlets, and FSC-A1/CFP+ for CFP+ cells. Cells were single-cell sorted using a SONY MA900 single-cell sorter with 300 eps max with an ultra purity setting into 96-well plates containing DMEM with 10% Tet-free FBS and Penn/strep. Uninduced infected U2OS 2-6-3 cells were used as a control for setting gates. CFP+ expression of each single cell clone was validated by flow cytometry on Guava (Cytek) after 500ng/mL doxycycline induction for 24 hours. Clones P24E and P32B were expanded and cultured in DMEM with 10% FBS and pen/strep.

### Validating Cas12a cleavage at HIV-ψ and 5’LTR enrichment with biotinylated DNA adapters

High molecular weight genomic DNA from clone P32B was used for enrichment analysis. As a no pulldown control, P32B HMW gDNA was digested with NcoI-HF and MspI (NEB) and purified. The pulldown library was made as follows: P32B HMW gDNA was digested with HIV-ψ-targeting Cas12a RNP, followed by heat inactivation at 65C. Biotinylated DNA adapters with HIV-PSI-R1 through HIV-PSI-R11 were ligated to the Cas12a digest directly in the Cas12a reaction mix with T4 DNA Ligase (NEB), followed by heat inactivation at 65C. The ligation was digested with MspI. Streptavidin pulldown was performed with the Dynabeads kilobaseBINDER (Invitrogen). Target DNA was released from the beads with NcoI-HF digest and purified. qPCR was performed with primers for mitochondrial DNA (non-target DNA) (mito_qpcr_F: GCCACAGCACTTAAACACATCTCT, mito_qpcr_R: TGAAATCTGGTTAGGCTGGTGTTAG), the Cas12a cut site within HIV-ψ (HIV_phi_F: ctctctcgacgcaggactc, HIV_phi_R: tttggcgtactcaccagtcg), and the 5’LTR (Target DNA) (5LTR_trunc_F: TCTGGCTAACTAGGGAACCCA, 5LTR_trunc_R: GGGCACACACTACTTGAAGC). **(Sup. Figure 3B).**

### Targeted Sequencing of Lentiviral Integration Sites from Bulk and Clonal Populations

Infected U2OS 2-6-3 cells from bulk or clonal populations (clone P32B or P24E) were cultured in DMEM with 10% FBS and pen/strep with 10uM BrdU or 10uM BrdU and 20ug/mL Puromycin for 24 hours. HMW gDNA was extracted and used to generate HIV-ψ nCasTLES libraries.

For the Bulk nCasTLES library, auxiliary DNA was generated by processing HMW gDNA from Bulk cells treated with BrdU-only for sequencing via Native Barcoding Kit 96 V14 (ONT). The Bulk BrdU-Only nCasTLES library was sequenced as follows: 1.64ng of the nCasTLES library was combined with 20.94ng of auxiliary DNA and sequenced on a MinION R10 Flow Cell (ONT) for 16 hours and 54 minutes. In a separate run, HMW gDNA from Bulk cells treated with 10uM BrdU and 20ug/mL Puromycin processed with the Native Barcoding Kit was sequenced on an R10 Flow Cell for 14 hours and 24 minutes. For the P24E nCasTLES libraries, auxiliary DNA was generated by processing gDNA from P24E cells treated with BrdU only or BrdU and Puromycin for sequencing via Native Barcoding Kit. The P24E BrdU-only nCasTLES library was sequenced as follows: 1.14ng of the nCasTLES was combined with 22.7ng of the P24E BrdU only auxiliary DNA library and sequenced on an R10 Flow Cell for 20 hours. The P24E BrdU/Puromycin nCasTLES library was sequenced as follows: 1.72ng of the nCasTLES was combined with 22.3ng of the P24E BrdU/Puromycin auxiliary DNA library and sequenced for 20 hours and 11 minutes. For the P32B nCasTLES library, carrier gDNA was prepared as follows: the high molecular weight gDNA from P32B cells treated with 10uM BrdU alone or with 10uM BrdU and 20ug/mL Puromycin were processed for sequencing via Native Barcoding Kit. 1.73ng of the nCasTLES library was combined with 23.25ng of the auxiliary DNA library and sequenced on an R10 Flow Cell for 51 hours.

Data processing was performed as described above, with the addition of the lentiviral 5’LTR and HIV-ψ to the reference. Reads containing the 5’LTR were extracted and re-aligned to hg38 to identify integration sites. 5mCG and BrdU calling were performed with Remora software from ONT using custom models. CNV analysis was performed with the R package QDNAseq^41^ and log2 ratios were plotted in R.

### Assessment of Cell Viability under Puromycin Treatment with CellTiter-Glo 2.0

Bulk and clonal populations (clone P24E and clone P32B) of infected U2OS 2-6-3 cells were cultured in DMEM with 10% FBS and pen/strep with puromycin (0ug/mL, 2ug/mL, 10ug/mL, 20ug/mL, 40ug/mL or 80ug/mL) for 24 hours. Viability was assessed with the ATP detection assay CellTiter-Glo 2.0.

---

**Supplemental Figure 1:**
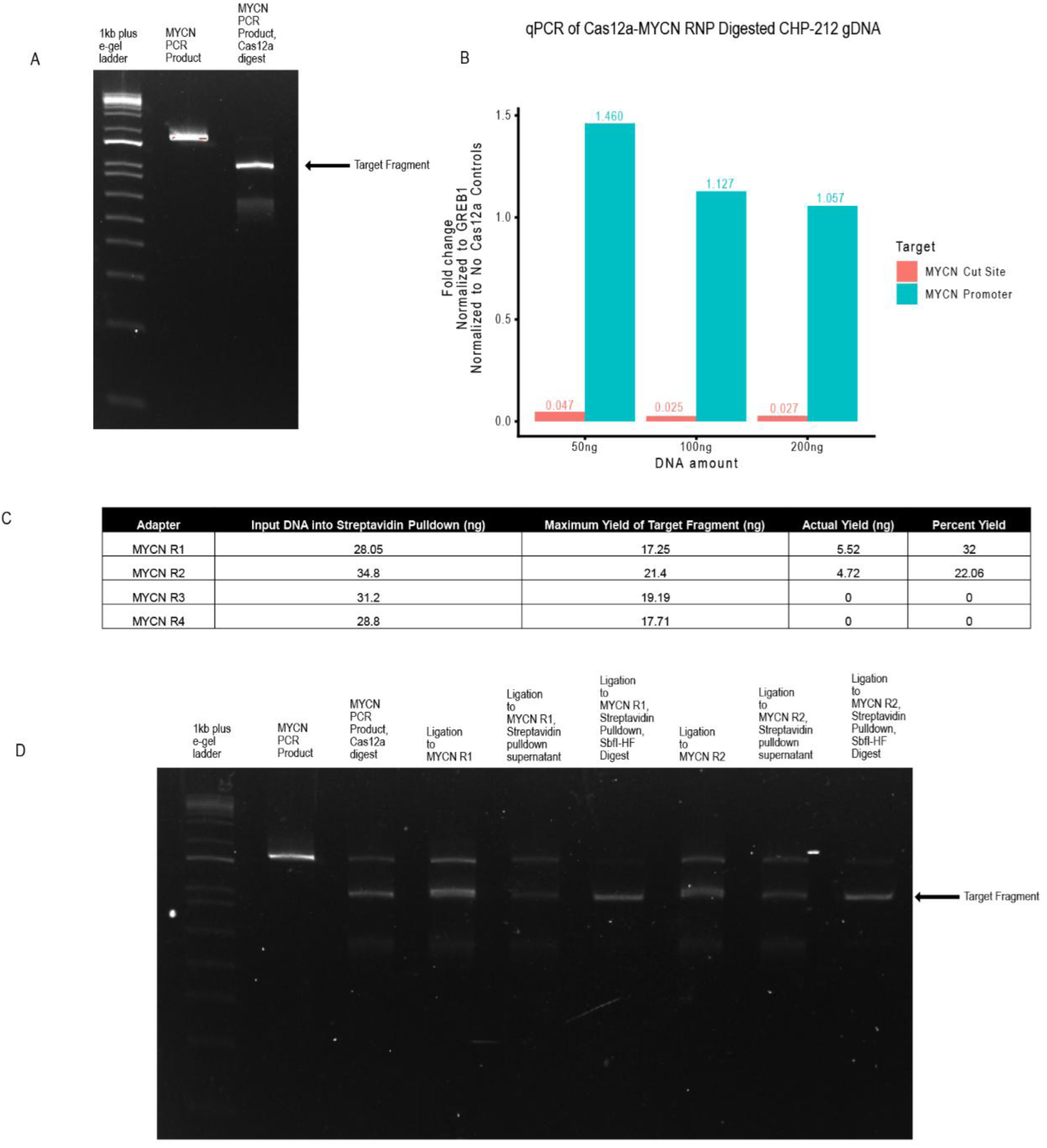
A. Agarose gel electrophoresis of MYCN promoter PCR product following cleavage with Cas12a-MYCN RNP. Lane 1 is the 1kb plus e-gel ladder, 2 is the full MYCN promoter PCR product and 3 is the MYCN promoter PCR product after Cas12a cleavage. B. Bar plot displays efficient on-target digest at the MYCN promoter with Cas12a-MYCN RNP with up to 200ng of input CHP-212 gDNA. C. Table displaying qubit quantification of MYCN promoter PCR product after streptavidin pulldown with biotinylated DNA adapters with 4 candidate 5’ overhangs. D. Agarose gel electrophoresis of MYCN promoter PCR product after streptavidin pulldown with biotinylated DNA adapters MYCN_R1 and MYCN_R2. Lane 1 is the 1kb plus e-gel ladder, Lane 2 is the full MYCN promoter PCR product, Lane 3 is the Cas12a-MYCN RNP digested PCR product, Lane 4 is the ligation to MYCN_R1, Lane 5 is the unbound fraction of the ligation to MYCN_R1 after streptavidin pulldown, Lane 6 is the captured on-target DNA of the ligation to MYCN_R1 after streptavidin pulldown and release with SbfI-HF digest, Lane 7 the ligation to MYCN_R2, Lane 8 is the unbound fraction of the ligation to MYCN_R2 after streptavidin pulldown, and Lane 9 is the captured on-target DNA of the ligation to MYCN_R2 after streptavidin pulldown and release with SbfI-HF digest.

**Supplemental Figure 2:**
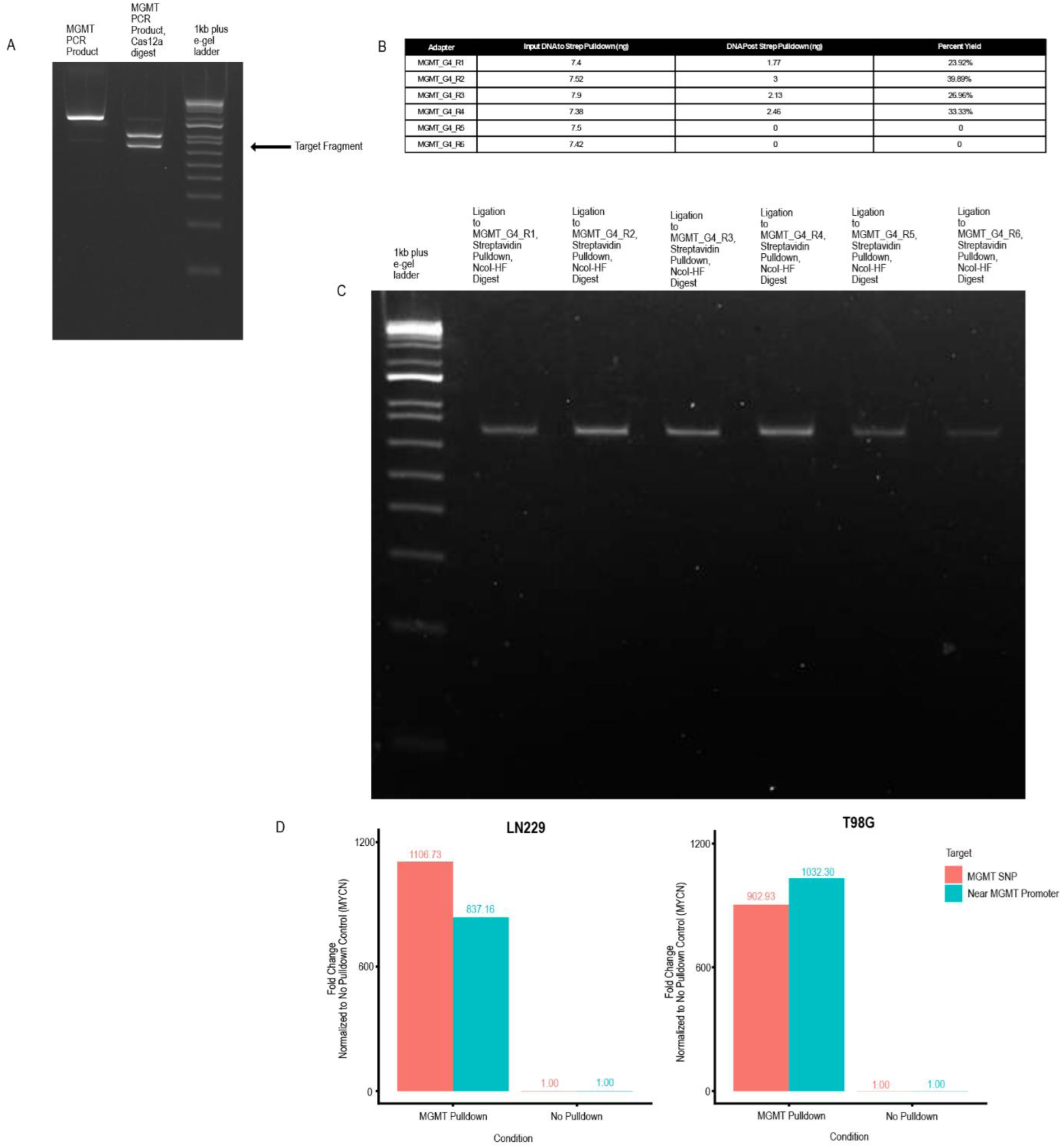
**A**. Agarose gel electrophoresis of MGMT promoter PCR product following cleavage with Cas12a-MGMT RNP. Lane 1 is the full MGMT promoter PCR product, 2 is the MGMT promoter PCR product after Cas12a cleavage, and 3 is the 1kb plus e-gel ladder. **B**. Table displaying qubit quantification of MYCN promoter PCR product after streptavidin pulldown with biotinylated DNA adapters with 6 candidate 5’ overhangs. Of the six predicted overhangs, four successfully ligated to a Cas12a-cleaved MGMT DNA amplicon and strongly enriched via streptavidin bead pulldown. **C**. Agarose gel electrophoresis of MGMT promoter PCR product after streptavidin pulldown with biotinylated DNA adapters MGMT_R1-6. **D.** Bar plot displays validation pulldown of MGMT promoter from LN229 or T98G gDNA with biotinylated DNA adapters with overhangs MGMT_R1-R4 via qPCR.

**Supplemental Figure 3.**
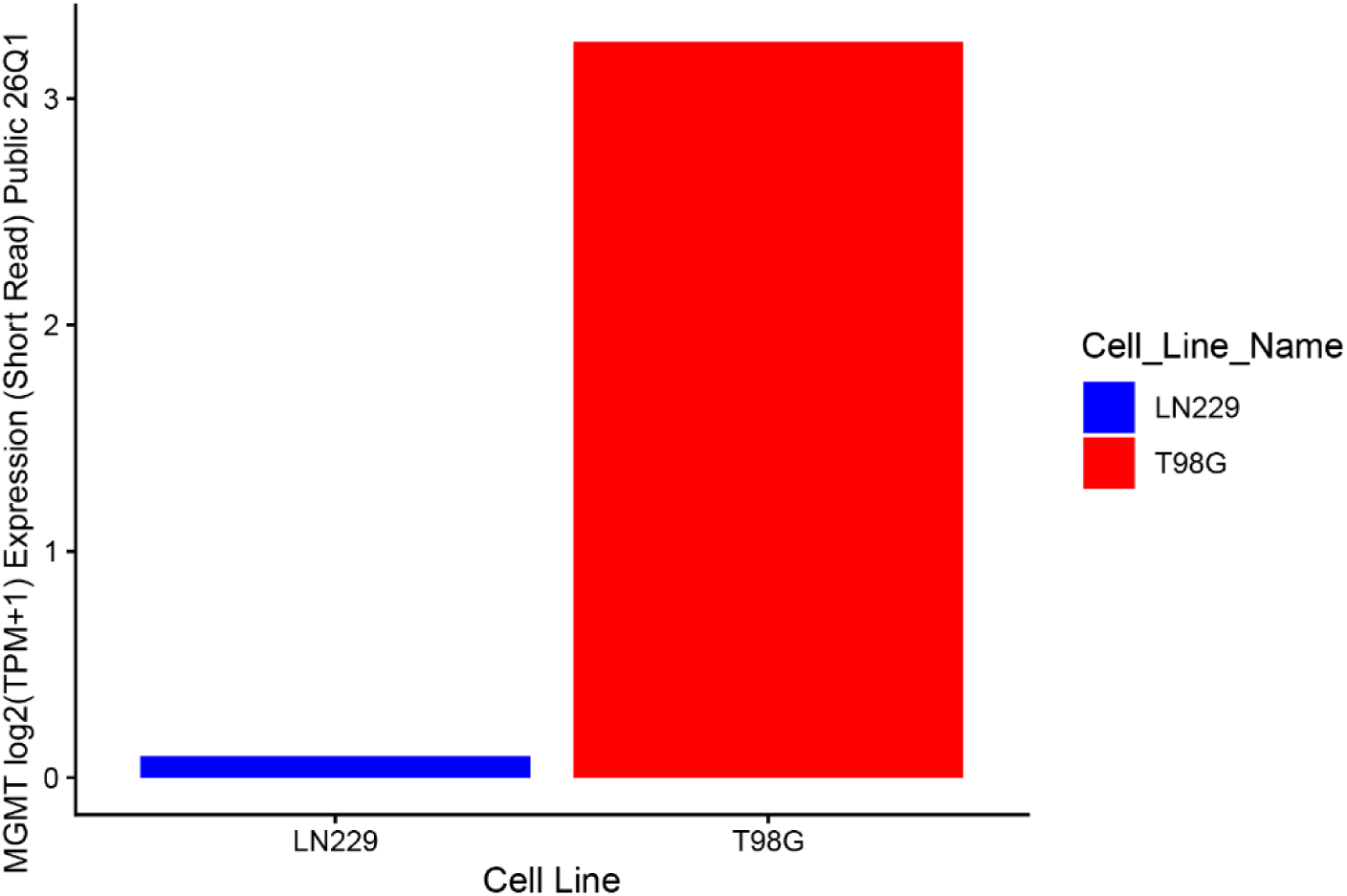
Expression of MGMT from LN229 and T98G cells, from depmap RNAseq data, plotted as log2(TPM+1).

**Supplemental Figure 4:**
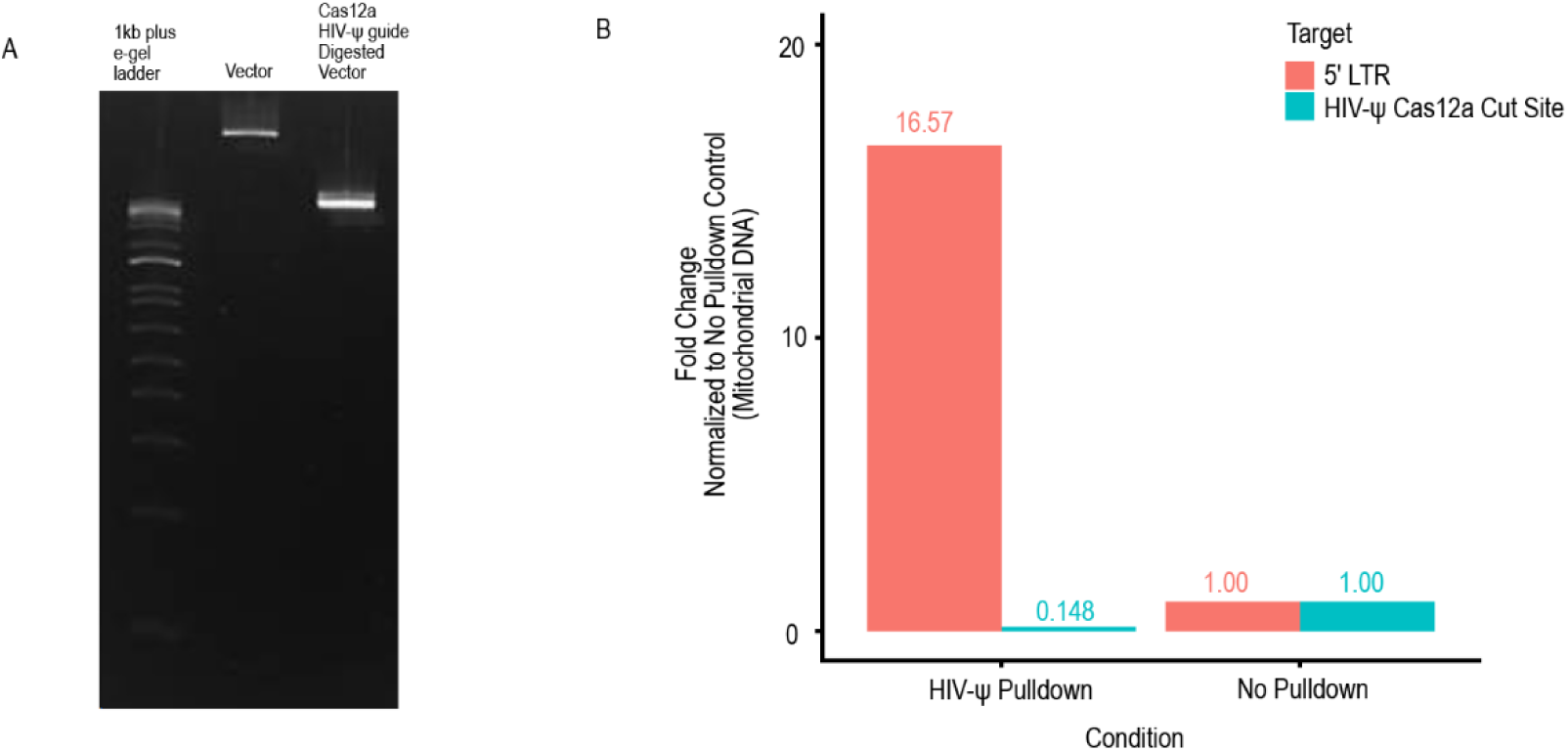
**A**. Agarose gel electrophoresis of pLentiCRISPRv1_SDHA following cleavage with Cas12a-HIV-ψ RNP. Lane 1 is the 1kb plus e-gel ladder, 2 is the pLentiCRISPRv1_SDHA and 3 is the linearized pLentiCRISPRv1_SDHA vector after Cas12a cleavage. **B**. Bar plot displays validation of on-target Cas12a-HIV-ψ RNP digest and of 5’LTR pulldown from U2OS 2-6-3 clone P32B gDNA with biotinylated DNA adapters with overhangs HIV-PSI-R1 to R11via qPCR.

**Supplemental Figure 5.**
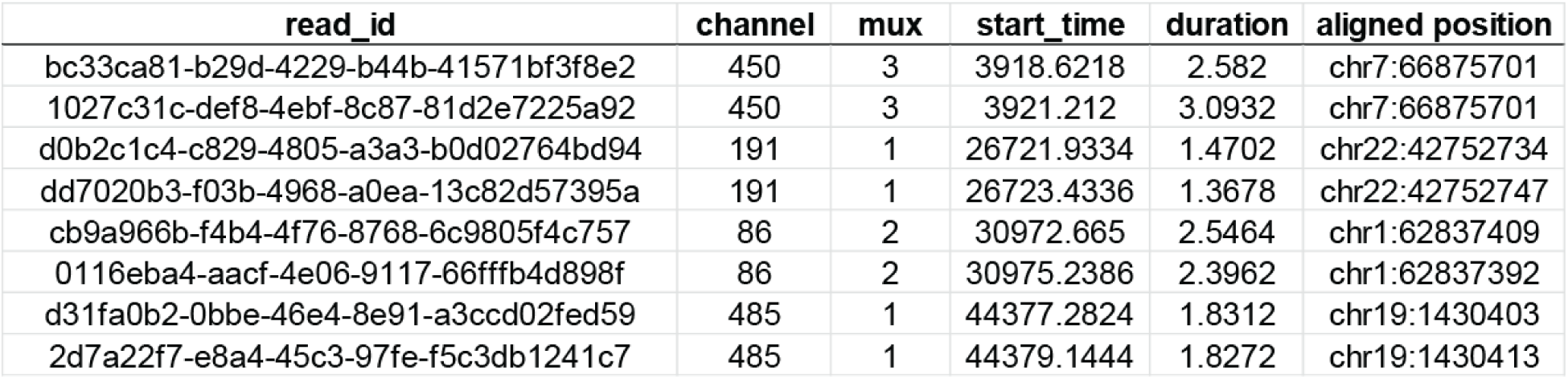
Table presents relevant information from sequencing summary file for each of the four pairs of captured HIVΨ reads that mapped to the same integration site. Channel refers to the pore cluster index and mux refers to the multiplexed pore within a specific channel, start_time and duration are in seconds, and genome positions are hg38. Notably, all four pairs of reads aligned to opposite strands of the reference and were sequenced sequentially through the exact same pore, and are thus duplexed reads representing both strands of a single captured DNA molecule.

**Supplemental Figure 6.**
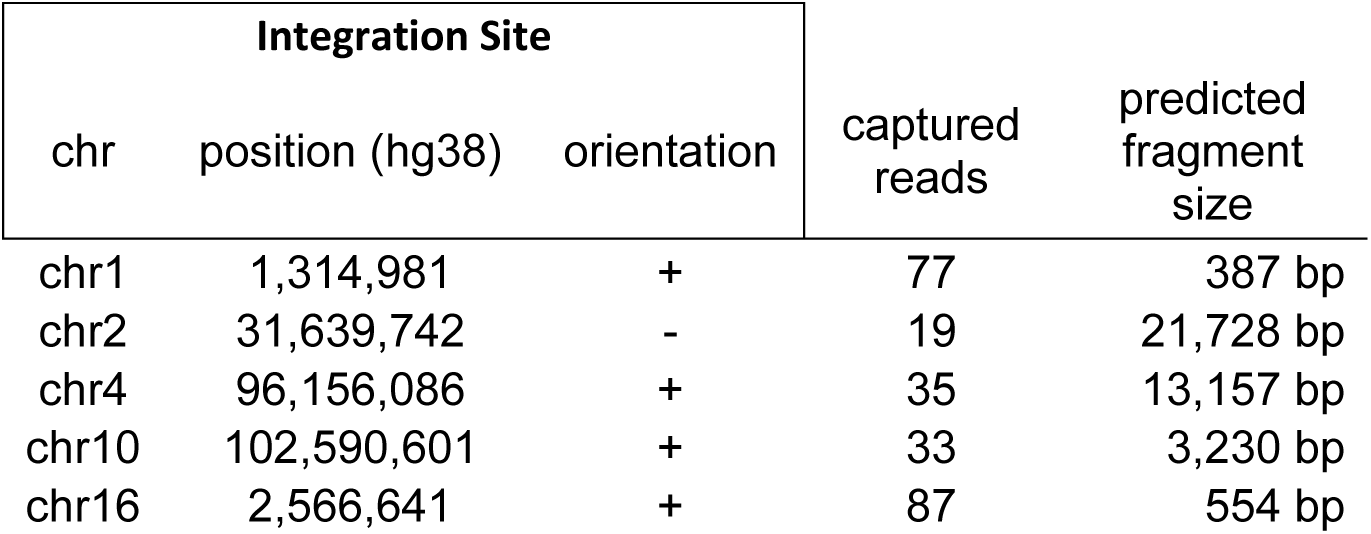
Table presents predicted fragment size of captured lentiviral fragments from clone P32B. Fragment size is predicted by distance from integration site to the nearest upstream MspI digest site, plus the expected fragment size of the captured lentiviral fragment.

