## Supplemental Note 1 nCasTLES protocol for "High-coverage DNA sequence and modification profiling of targeted genomic elements using Nanopore-based Cas12a Targeted Ligation and Enrichment Sequencing (nCasTLES)"

**Contents:**

Supplementary Note 1. Detailed Cas12a-mediated Target Enrichment for Nanopore Sequencing Protocol

**Supplemental Note 1, Cas12a-mediated Target Enrichment for Nanopore Sequencing Protocol**

Date: 07/16/2026

Author: Milo Vantine

Notes: In this protocol, I will describe the workflow for performing target enrichment using Cas12a for Nanopore sequencing of native DNA. I will use target enrichment of the MGMT promoter from the LN229 cell line as an example. Cas12a guides, biotin adapters and PCR primers should be customized for targets of interest.

Overview:

Step 1: High Molecular Weight gDNA Extraction

Step 2: Determine 4- and 5- base overhangs generated by Cas12a digest of target

Step 3: Cas12a-mediated Target Enrichment for Nanopore Sequencing

Step 4: Prepare carrier DNA

Step 5: Sequence

Materials:

Table 1: Oligonucleotides

**Oligos**

| Type | Example Oligo Name | Example Sequence | Modifications |
|---|---|---|---|
| Cas12a guide RNA | MGMT_Guide_4 | rUrArArUrUrUrCrUrArCrUrArArGrUrGrUrArGrArUrGrUrGrUrCrCrUrArArGrArArArGrGrGrUrGrGrGrArArG | None |
| PCR Adapter F | Amp_seq_F | TGGAAAGGATTCATTCCCACGGCTTGGGATGAATGGCAACTGTCGAGGACCTATCCACTAGACGAGTATTGCTGCAGTCTGCA | None |
| PCR Adapter 4N R | Amp_seq_4N_R | /5Phos/NNNNTGCAGACTGCAGCAATACTCGTCTAGTGGATAGGTCCTCGACAGTTGCCATTCATCCCAAGCCGTGGGAATGAATCCTTTCCA | 5' Phosphorylated |
| PCR Adapter 5N R | Amp_seq_5N_R | /5Phos/NNNNNTGCAGACTGCAGCAATACTCGTCTAGTGGATAGGTCCTCGACAGTTGCCATTCATCCCAAGCCGTGGGAATGAATCCTTTCCA | 5' Phosphorylated |
| PCR Primer (for PCR adapter) | bc7_qPCR_R | GGAAAGGATTCATTCCCACGG | None |
| Locus Specific PCR Primer | qPCR_MGMT_promoter_enrich_F | AGGGCTCATCCCAAAGCAAA | None |
| Biotin Adapter F | SbfI_biotin_TEG_F | /5BiotinTEG/TGGCCTAGGGAGCTCCCATGGTGATCAGGATCCCCTGCAGGTCTTGG | 5' Biotin TEG |
| Biotin Adapter R | MGMT_G4_R1 | /5Phos/CTTCCCAAGACCTGCAGGGGATCCTGATCACCATGGGAGCTCCCTAGGCCAC | 5' Phosphorylated |
| Biotin Adapter R | MGMT_G4_R2 | /5Phos/TTCCCCAAGACCTGCAGGGGATCCTGATCACCATGGGAGCTCCCTAGGCCAC | 5' Phosphorylated |
| Biotin Adapter R | MGMT_G4_R3 | /5Phos/TCCCCCAAGACCTGCAGGGGATCCTGATCACCATGGGAGCTCCCTAGGCCAC | 5' Phosphorylated |
| Biotin Adapter R | MGMT_G4_R4 | /5Phos/CTTCCCCAAGACCTGCAGGGGATCCTGATCACCATGGGAGCTCCCTAGGCCAC | 5' Phosphorylated |

Table 2: Buffers, Enzymes and other Reagents

| Name | Supplier | Catalog Number | Type | Recipe | Step |
|---|---|---|---|---|---|
| Isopropanol | Sigma-Aldrich | I9516-25M | Alcohol | N/A | High Molecular Weight gDNA Extraction |
| Corning™ Cell Culture Phosphate Buffered Saline (1X) | Corning | 21040CV | Buffer | N/A | High Molecular Weight gDNA Extraction |
| Annealing Buffer | N/A | N/A | Buffer | 100mM Tris-HCl pH 7.5, 2M LiCl | All |
| NEBuffer™ r2.1 | New England Biolabs | B6002S | Buffer | N/A | Amplicon Sequencing, Cas12a Enrichment |
| EB Buffer | N/A | N/A | Buffer | 10mM Tris-HCl pH 8.0 | Amplicon Sequencing, Cas12a Enrichment |
| Q5 Reaction Buffer (5x) | New England Biolabs | B9027S | Buffer | N/A | Amplicon Sequencing |
| rCutSmart™ Buffer (10X) | New England Biolabs | B6004S | Buffer | N/A | Cas12a Enrichment |
| Triton™ X-100 | Sigma-Aldrich | X100-5ML | Detergent | N/A | High Molecular Weight gDNA Extraction |
| Proteinase K, Molecular Biology Grade | New England Biolabs | P8107S | Enzyme | N/A | High Molecular Weight gDNA Extraction |
| Monarch® RNase A | New England Biolabs | T3018L | Enzyme | N/A | High Molecular Weight gDNA Extraction |
| EnGen® Lba Cas12a (Cpf1) | New England Biolabs | M0653S | Enzyme | N/A | Amplicon Sequencing, Cas12a Enrichment |
| T4 DNA Ligase | New England Biolabs | M0202L | Enzyme | N/A | Amplicon Sequencing, Cas12a Enrichment |
| Q5® High-Fidelity DNA Polymerase | New England Biolabs | M0491S | Enzyme | N/A | Amplicon Sequencing |
| NcoI-HF | New England Biolabs | R3193S | Enzyme | N/A | Cas12a Enrichment |
| PflMI | New England Biolabs | R0509S | Enzyme | N/A | Cas12a Enrichment |
| Salt-T4 DNA Ligase | New England Biolabs | M0467S | Enzyme | N/A | Cas12a Enrichment |
| Blunt/TA Ligase Master Mix | New England Biolabs | M0367S | Enzyme | N/A | Cas12a Enrichment |
| Monarch® HMW DNA Extraction Kit for Cells & Blood | New England Biolabs | T3050L | Kit | N/A | High Molecular Weight gDNA Extraction |
| Qubit™ dsDNA HS Quantification Assay | Invitrogen | Q32851 | Kit | N/A | All |
| Monarch® Spin PCR & DNA Cleanup Kit (5 μg) | New England Biolabs | T1130S | Kit | N/A | Amplicon Sequencing, Cas12a Enrichment |
| Native Barcoding Kit 24 V14 | Oxford Nanopore Technologies | SQK-NBD114.24 | Kit | N/A | Cas12a Enrichment |
| Ligation Sequencing Kit V14 | Oxford Nanopore Technologies | SQK-LSK114 | Kit | N/A | Cas12a Enrichment |
| Dynabeads™ kilobaseBINDER™ Kit | Invitrogen | 60101 | Kit | N/A | Cas12a Enrichment |
| NEBNext® Ultra™ II End Repair/dA-Tailing Module | New England Biolabs | E7546S | Kit | N/A | Cas12a Enrichment |
| Adenosine 5'-Triphosphate (ATP) (10mM) | New England Biolabs | P0756S | Nucleotide | N/A | Amplicon Sequencing, Cas12a Enrichment |
| Deoxynucleotide (dNTP) Solution Mix (10mM) | New England Biolabs | N0447S | Nucleotide | N/A | Amplicon Sequencing |
| AMPure XP Beads for DNA Cleanup, 5 mL | Beckman Coulter | A63880 | Spri Reagent | N/A | Amplicon Sequencing, Cas12a Enrichment |

Table 3: Other Materials

| Name | Supplier | Catalog Number |
| --- | --- | --- |
| ART™ Wide Bore Filtered Pipette Tips (1000uL) | Thermo Scientific | 2079GPK |
| ART™ Wide Bore Filtered Pipette Tips (200uL) | Thermo Scientific | 2069GPK |
| DynaMag™-2 Magnet | Invitrogen | 12321D |
| Eppendorf™ DNA LoBind™ Tubes (1.5mL) | Eppendorf | 30108418 |
| Eppendorf™ DNA LoBind™ Tubes (2mL) | Eppendorf | 22431048 |
| MinION/GridION Flow Cell (DNA) | Oxford Nanopore Technologies | FLO-MIN114 |

Protocol:

*Step 1: High Molecular Weight gDNA Extraction (2 Days)*

1. Start with a cell pellet containing 10^6 live cells (fresh or flash frozen)
2. Resuspend the cell pellet in 100uL of ice-cold PBS

The steps below utilize the Monarch® HMW DNA Extraction Kit for Cells & Blood kit (NEB, T3050L):

3. Make enough EBII+ buffer (EBII Buffer, 0.02% Triton-X 100) so that you have 200uL per sample
4. Aliquot 200uL of EBII+ buffer into labeled 2mL Lobind Eppendorf tubes. These will be used later in the protocol for elution of the gDNA from DNA Capture Beads. Set the tubes aside at RT.
5. For each sample, create a premix of Direct Lysis Mix by combining the following in a 2mL Lo-bind Eppendorf tube:
    a. 100 µl Nuclei Prep Buffer
    b. 100 µl Nuclei Lysis Buffer
    c. 10 µl Proteinase K
6. Add 200 µl Direct Lysis Mix to resuspended cells and mix by inverting 10 times.
7. Incubate for 10 minutes at 56°C with thermal mixer set at 600 RPM
8. Add 5uL of RNAse A to each sample, and pipette up and down gently 10 times with a wide-bore pipette tip
9. Incubate at 56C on a heat block for 10 minutes
10. Add 75uL of Precipitation Enhancer to each sample and mix by inverting the tube 10 times
11. Add 2 DNA Capture Beads to each sample
12. Add 275 µl isopropanol to each sample.
13. Invert the sample via slow manual inversion 25 times (~5 seconds per full inversion).
14. Remove and discard the liquid by pipetting. Try not to touch the beads with your pipette tip. If necessary, leave some liquid behind.
15. Add 500 µl gDNA Wash Buffer, close the cap, and mix by inverting the tube 2–3 times.
16. Remove and discard the liquid by pipetting.
17. Add 500 µl gDNA Wash Buffer, close the cap, and mix by inverting the tube 2–3 times.
18. Remove and discard the liquid by pipetting.
19. Place a bead retainer in a 2mL collection tube for each sample
20. Carefully pour the beads into the bead retainer and close the cap
21. Tap the bead retainer gently on absorbent paper to remove traces of wash buffer.
22. Pour the beads into the 2mL tubes containing 200uL of EBII+ buffer
23. Incubate the tubes at 56°C on a heat block to elute the DNA off the beads. To facilitate elution during incubation, gently pipette and dispense the eluate a few times over the glass beads using a wide bore pipette tip.
24. Incubate the tubes at RT overnight.
25. Place a bead retainer in a labeled 1.5mL Lobind Eppendorf tube for each sample
26. Pour the beads and elution buffer into the bead retainer.
27. Centrifuge for 1 minute at maximum speed (>16,000 x g). Check if DNA was completely released from the beads. If after the spin DNA threads are visible between beads and eluate, centrifuge for 1 additional minute at max speed.
28. Remove the bead retainer and beads and discard.
29. Using a wide bore pipette tip, carefully pipette eluates up and down 5 times for homogenization.

30. Incubate the eluates at 37°C on a heat block for 10 minutes.
31. Repeat steps 29 and 30, if needed.
32. NOTE: At this stage, the eluates are usually fairly sticky, which can make quantification challenging. However, the stickiness of these eluates does not seem to hinder downstream processing.
33. Quantify eluates using the Qubit™ dsDNA HS Quantification Assay
34. Store eluates at 4C short-term and -20C for long-term storage.

*Step 2: Determine 4- and 5- base overhangs generated by Cas12a digest of target*

Note: Reactions throughout this protocol can be scaled up or down, depending on target of interest. Input DNA can be gDNA as in the example below. If input DNA is a PCR product or a vector containing the region of interest, the reactions can be scaled down.

1. <u>Anneal 4N- and 5N- overhang PCR adapters as follows:</u>
    a. In PCR tubes, set the following:

| 4N Adapter Annealing Reaction | |
|---|---|
| **Reagent** | **Volume (uL)** |
| Annealing Buffer | 1 |
| Amp_seq_F (100uM) | 1 |
| Amp_seq_4N_R (100uM) | 1 |
| Water | 2.5 |

| 5N Adapter Annealing Reaction | |
|---|---|
| **Reagent** | **Volume (uL)** |
| Annealing Buffer | 1 |
| Amp_seq_F (100uM) | 1 |
| Amp_seq_5N_R (100uM) | 1 |
| Water | 2.5 |

    b. Incubate annealing reactions at 95°C for 5 minutes in a thermal cycler with a heated lid
    c. Incubate the annealing reactions at RT on the benchtop for 15 minutes
2. <u>Digest target gDNA with Cas12a as follows:</u>
    a. Determine the volume of gDNA needed for 4.5ug
        i. In this example, I use 96uL of LN229 HMW gDNA
    b. Assemble the Cas12a RNPs by combining the following in a 1.5mL tube:

| Cas12a RNP Mix | |
|---|---|
| **Reagent** | **Vol (uL)** |
| NEBuffer R2.1 (10X) | 24 |
| MGMT_Guide_4 (1uM) | 8 |
| Lba Cas12a (1uM) | 8 |
| Water | 104 |

        i. Note: For these reactions, add water to a total volume of 240uL – volume of gDNA to add later
            1. EX: I will use 96uL of gDNA in this reaction, so I add water to a total volume of 144uL in this reaction.
    c. Incubate the Cas12a RNP mix at 25°C for 10 minutes
    d. Add 4.5ug of target DNA to the Cas12a RNP Mix
        i. In this example, I added 96uL of LN229 HMW gDNA
    e. Incubate reaction at 37°C on the heat block for 15 minutes
    f. Incubate the reaction at 65°C on the heat block for 15 minutes to inactivate the Cas12a
3. <u>Ligate 4N- and 5N- PCR Adapters as follows:</u>
    a. Set the following ligation reactions in 1.5mL Lobind Eppendorf tubes:

| 4N- Adapter Ligation Reaction | |
|---|---|
| **Reagent** | **Vol (uL)** |
| NEBuffer r2.1 (10X) | 12 |
| ATP (10mM) | 24 |
| Cas12a Reaction | 120 |
| 4N (annealed) | 1 |
| T4 DNA Ligase | 12 |
| Nuclease-free Water | 71 |

| 5N- Adapter Ligation Reaction | |
|---|---|
| **Reagent** | **Vol (uL)** |
| NEBuffer r2.1 (10X) | 12 |
| ATP (10mM) | 24 |
| Cas12a Reaction | 120 |
| 5N (annealed) | 1 |
| T4 DNA Ligase | 12 |
| Nuclease-free Water | 71 |

    b. Incubate ligation reactions at RT on the benchtop for 10 minutes
    c. Clean up the reactions with 0.4X Ampure XP size selection to remove extra adapters as follows:
        i. Add 96uL of Ampure XP beads to each reaction

ii. Flick the tubes to mix well
iii. Incubate tubes on a Hula Mixer at RT for 10 minutes
iv. Place tubes on a magnetic rack to pellet the beads
v. Remove and discard the supernatant without disturbing the beads
vi. Keeping the tubes on the magnetic rack, wash the beads twice with fresh 85% ethanol
vii. Remove and discard the ethanol
viii. Briefly spin down the tubes in a microcentrifuge
ix. Place tubes back on the magnetic rack to pellet the beads
x. Remove residual ethanol
xi. Allow the pellets to air dry for 30 seconds
xii. Resuspend each pellet in 130uL of EB Buffer
xiii. Incubate the tubes at 37°C on a heat block for 15 minutes, flicking the tubes every 5 minutes
xiv. Place tubes back on the magnetic rack to pellet the beads
xv. Transfer the eluates to labeled 1.5mL Lobind Eppendorf tubes

d. Quantify eluates using the Qubit™ dsDNA HS Quantification Assay

4. Perform PCR to amplify over the Cas12a cut site as follows:
   a. Set 12 each of the following PCR reactions in PCR tubes:

| 4N Overhang PCR | |
|---|---|
| **Reagent** | **Vol (uL)** |
| Q5 Reaction Buffer (5X) | 10 |
| 4N Adapter Ligation Reaction | 10 |
| dNTPs (10mM) | 1 |
| MGMT_G4_Amp_F (10uM) | 2.5 |
| bc7_qPCR_R (10uM) | 2.5 |
| Q5 Polymerase | 0.5 |
| Water | 23.5 |

| 5N Overhang PCR | |
|---|---|
| **Reagent** | **Vol (uL)** |
| Q5 Reaction Buffer (5X) | 10 |
| 5N Adapter Ligation Reaction | 10 |
| dNTPs (10mM) | 1 |
| MGMT_G4_Amp_F (10uM) | 2.5 |
| bc7_qPCR_R (10uM) | 2.5 |
| Q5 Polymerase | 0.5 |
| Water | 23.5 |

   b. Cycle PCR reactions as follows in a thermal cycler:

| STEP | TEMP | TIME |
|---|---|---|
| Initial Denaturation | 98°C | 30 seconds |
| 40 Cycles | 98°C | 5 sec |
| | 66C | 10 sec |
| | 72°C | 30 sec |
| Final Extension | 72°C | 2 minutes |
| Hold | 4°C | |

   c. Clean up reactions following the manufacturer's guidelines for the Monarch® Spin PCR & DNA Cleanup Kit (5 µg) (NEB). Purify 3 PCR reactions per column and elute from each column with 20uL of EY Buffer. Combine the like eluates into 1.5mL tubes (80uL total volume per sample).
   d. Optional: Perform additional size selection with an Ampure XP cleanup. For this example, I performed a 0.6X Ampure XP size selection as follows:
      i. Add 48uL of Ampure XP beads to each reaction
      ii. Flick the tubes to mix well
      iii. Incubate tubes on a Hula Mixer at RT for 10 minutes
      iv. Place tubes on a magnetic rack to pellet the beads
      v. Remove and discard the supernatant without disturbing the beads
      vi. Keeping the tubes on the magnetic rack, wash the beads twice with fresh 85% ethanol
      vii. Remove and discard the ethanol
      viii. Briefly spin down the tubes in a microcentrifuge
      ix. Place tubes back on the magnetic rack to pellet the beads
      x. Remove residual ethanol
      xi. Allow the pellets to air dry for 30 seconds

xii. Resuspend each pellet in 15uL of EB Buffer
xiii. Incubate the tubes at 37°C on a heat block for 15 minutes, flicking the tubes every 5 minutes
xiv. Place tubes back on the magnetic rack to pellet the beads
xv. Transfer the eluates to labeled 1.5mL Lobind Eppendorf tubes

e. Quantify eluates using the Qubit™ dsDNA HS Quantification Assay
f. Run final PCR product on an agarose gel to verify the size.
g. Submit to Genewiz for PCR-EZ (Long read amplicon sequencing).
h. Use the PCR-EZ results to determine the 4- and 5-base 5’ overhangs left at your region of interest by Cas12a.

*Step 3: Nanopore Cas-12a Targeted Ligation-Enrichment Sequencing (nCasTLES)*

1. Anneal biotin adapter oligos as follows:
   a. Set one annealing reaction for each biotin adapter as follows:

| Biotin Adapter 1 | |
|---|---|
| **Reagent** | **Vol (uL)** |
| Milo's Annealing Buffer | 1 |
| SbfI_biotin_TEG_F (100uM) | 1 |
| MGMT_G4_R1 (100uM) | 1 |
| Water | 2.5 |

| Biotin Adapter 2 | |
|---|---|
| **Reagent** | **Vol (uL)** |
| Milo's Annealing Buffer | 1 |
| SbfI_biotin_TEG_F (100uM) | 1 |
| MGMT_G4_R2 (100uM) | 1 |
| Water | 2.5 |

| Biotin Adapter 3 | |
|---|---|
| **Reagent** | **Vol (uL)** |
| Milo's Annealing Buffer | 1 |
| SbfI_biotin_TEG_F (100uM) | 1 |
| MGMT_G4_R3 (100uM) | 1 |
| Water | 2.5 |

| Biotin Adapter 4 | |
|---|---|
| **Reagent** | **Vol (uL)** |
| Milo's Annealing Buffer | 1 |
| SbfI_biotin_TEG_F (100uM) | 1 |
| MGMT_G4_R4 (100uM) | 1 |
| Water | 2.5 |

      i. In this example, we had 4 different overhangs left by the Cas12a digest at the MGMT locus. Each annealing reaction contains the biotinylated universal forward oligo and a reverse oligo containing a 4- or 5- base overhang compatible with each Cas12a cut. Some targets will require more or fewer adapters.
   b. Incubate annealing reactions at 95°C for 5 minutes in a thermal cycler with a heated lid
   c. Incubate the annealing reactions at RT on the benchtop for 15 minutes
   d. Add 10uL of nuclease-free water to each annealing reaction and quantify using the Qubit™ dsDNA HS Quantification Assay.
   e. For the ligation reactions, aim for 500ng total of biotin adapter oligo. For this example, that means we will aim for 125ng of each biotin adapter.
      i. In a case where more adapters are required, scale down the mass of each individual adapter, aiming for a total of 500ng. For example, if you are using 10 adapters, use 50ng of each adapter in the ligation reaction.
2. Digest target gDNA with Cas12a as follows:
   a. Determine the volume of gDNA needed for 2.5ug
      i. In this example, I use 85uL of T98G HMW gDNA
   b. Assemble the Cas12a RNPs by combining the following in a 1.5mL tube:

| Cas12a RNP Mix | |
|---|---|
| **Reagent** | **Vol (uL)** |
| NEBuffer R2.1 (10X) | 60 |
| MGMT_Guide_4 (1uM) | 20 |
| Lba Cas12a (1uM) | 20 |
| Water | 415 |

      i. Note: For these reactions, add water to a total volume of 600uL – volume of gDNA to add later
         1. EX: I will use 85uL of T98G gDNA in this reaction, so I add water to a total volume of 415uL in this reaction.
   c. Incubate the Cas12a RNP mix at 25°C for 10 minutes
   d. Add >1ug* of target DNA to the Cas12a RNP Mix

        i. Different targets will require different amounts of input. For high copy number targets, 1ug of input DNA may be sufficient. For low copy number targets, 2-5ug of input gDNA is recommended.
        ii. In this example, 2.5ug (85uL) of T98G gDNA was used as the input.
    e. Incubate reaction at 37°C on the heat block for 15 minutes
    f. Incubate the reaction at 65°C on the heat block for 15 minutes to inactivate the Cas12a
    g. Place tubes briefly on ice to cool before proceeding to ligation reactions
3. <u>Ligate biotin adapters as follows:</u>
    a. Set the following ligation reactions in 1.5mL tubes:

| Biotin Adapter Ligation Reaction | |
|---|---|
| **Reagent** | **Vol (uL)** |
| NEBuffer r2.1 (10X) | 20 |
| ATP (10mM) | 80 |
| Cas12a Reaction | 600 |
| Biotin Adapter 1 | 125ng |
| Biotin Adapter 2 | 125ng |
| Biotin Adapter 3 | 125ng |
| Biotin Adapter 4 | 125ng |
| T4 DNA Ligase | 40 |
| Nuclease-free Water | To a total volume of 800uL |

    b. Incubate at RT for 10 minutes on the benchtop
    c. Heat inactivate at 65C for 15 minutes on the heat block
    d. Briefly place tubes on ice to cool down before proceeding to next step
4. <u>Digest your ligation reaction with restriction enzyme of your choice as follows:</u>
    a. In this example, we used PflMI for the restriction digests.
    b. Add 16uL of PflMI to the ligation reaction mix.
    c. Incubate at 37C on the heat block for 15 minutes
    d. Incubate at 65C on the heat block for 20 minutes
    e. Briefly place tubes on ice to cool down before proceeding to next step
5. <u>Enrich for target of interest with streptavidin pulldown as follows:</u>
    a. Set 100uL of Dynabeads™ kilobaseBINDER™ beads in one 1.5mL (per sample)
    b. Place tube on magnetic rack for 2 minutes
    c. Remove and discard the supernatant
    d. Gently resuspend the beads in 400uL Binding Solution
    e. Place tube on magnetic rack for 2 minutes
    f. Remove and discard the supernatant
    g. Gently resuspend the beads in 816uL Binding Solution
    h. Add 816uL of ligation reaction to each tube using a wide-bore tip
    i. Gently pipette with a wide bore tip to mix
    j. Incubate at RT for 3 hours on the Hula mixer at RT
    k. Place tube on magnetic rack for 2 minutes
    l. Remove and discard the supernatant
    m. Wash the Dynabeads®/DNA-complex twice in 800μL Washing Solution and once in water
        i. Resuspend the beads each time. Incubate on the magnetic rack for 2 minutes each time.
    n. To release target DNA from the beads, we will perform a restriction digest that cleaves the Biotin-TEG tag from the biotin adapter. Resuspend the beads in each tube each in:
        i. 220uL water
        ii. 25uL 10X rCutsmart Buffer
        iii. 5uL NcoI-HF (Or enzyme of your choice)
    o. Incubate restriction digest on the heat block at 37C for 15 minutes, gently flicking the tube every few minutes to make sure the beads don't settle to the bottom
    p. Place tube back on the magnetic rack for 2 minutes to pellet the beads
    q. Transfer the eluate to a labeled 1.5mL Lobind Eppendorf tube
    r. Clean up the eluate following the manufacturer's guidelines for the Monarch® Spin PCR & DNA Cleanup Kit (5 μg) (NEB).

s. Elute from the monarch column with 25uL warm EY buffer
t. Quantify eluates using the Qubit™ dsDNA HS Quantification Assay
u. Remove excess adapter fragments with 0.9X Ampure XP cleanup as follows:
    i. Added 22.5uL ampure XP beads to the sample
    ii. Placed on hula mixer at RT for 5 minutes
    iii. Place tube on magnetic rack for 5 minutes
    iv. Remove and set aside supernatant (just in case)
    v. Wash the beads twice with 200uL of 80% ethanol, keeping the tube on the magnetic rack
    vi. Air dry the pellet on the magnetic rack for up to 1 minute
    vii. Resuspend the ampure XP beads in 48uL EB buffer
    viii. Incubate on the heat block at 37C for 15 minutes, flicking the tubes occasionally
    ix. Place tube on the magnetic rack to pellet the beads
    x. Transfer eluate to a new labeled 1.5mL tube
    xi. NOTE: You may choose to qubit quant your library here. You **should** expect an extremely low quant or for your DNA concentration to be **too low to quantify.**
v. Store Cas12a enrichment library at -20C long term or proceed to next step

6. Prepare Cas12a enrichment library for sequencing with the Ligation Sequencing Kit V14 (Oxford Nanopore Technologies):
    a. Thaw the following on ice:
        i. NEBNext® FFPE DNA Repair Mix (M6630)
        ii. NE Next® FFPE DNA Repair Buffer v2 (E7363)
        iii. NEBNext® Ultra II End Prep Enzyme Mix (E7646)
    b. Vortex the FFPE DNA Repair Buffer v2 to ensure it is well mixed.
    c. Set the following in a PCR tube:

| Reagent | Vol (uL) |
|---|---|
| Cas12a enrichment library | 47 |
| NEBNext FFPE DNA Repair Buffer v2 | 7 |
| NEBNext FFPE DNA Repair Mix | 2 |
| Ultra II End-prep Enzyme Mix | 3 |

    d. Thoroughly mix the reaction by gently pipetting and briefly spinning down.
    e. Using a thermal cycler, incubate at 20°C for 5 minutes and 65°C for 5 minutes. Then cool down to between 4°C and 20°C on the thermal cycler or place the samples on ice.
    f. Resuspend the AMPure XP Beads by vortexing.
    g. Spin down and transfer the DNA sample to a clean 1.5 ml Eppendorf DNA LoBind tube.
    h. Add 60 μl of resuspended the AMPure XP Beads to the end-prep reaction and mix by flicking the tube
    i. Incubate on a Hula mixer for 15 minutes at room temperature
    j. Prepare 500 μl of fresh 80% ethanol in nuclease-free water.
    k. Spin down the sample and pellet on a magnet until supernatant is clear and colourless. Keep the tube on the magnet, and pipette off the supernatant.
    l. Keep the tube on the magnet and wash the beads with 200 μl of freshly prepared 80% ethanol without disturbing the pellet. Remove the ethanol using a pipette and discard.
    m. Repeat the previous step.
    n. Spin down and place the tube back on the magnet. Pipette off any residual ethanol. Allow to dry for ~30 seconds, but do not dry the pellet to the point of cracking.
    o. Remove the tube from the magnetic rack and resuspend the pellet in 30 μl nuclease-free water. Incubate for 10 minutes at 37C
    p. Pellet the beads on a magnet until the eluate is clear and colourless, for at least 1 minute.
    q. Remove and retain 30 μl of eluate into a clean 1.5 ml Eppendorf DNA LoBind tube.
    r. Resuspend in 31uL of nuclease free water. Incubate at 37C for 10 minutes
    s. Pellet the beads on a magnet until the eluate is clear and colourless, for at least 1 minute.
    t. Move the 31uL of eluate to the 1.5mL tube with the first 30uL of eluate (61uL total)
    u. Quantify eluate using the Qubit™ dsDNA HS Quantification Assay
    v. Spin down the Ligation Adapter (LA) and Salt T4 DNA Ligase, and place on ice.

w. Thaw Ligation Buffer (LNB) at room temperature, spin down and mix by pipetting. Due to viscosity, vortexing this buffer is ineffective. Place on ice immediately after thawing and mixing.
x. Thaw the Elution Buffer (EB) at room temperature and mix by vortexing. Then spin down and place on ice
y. Thaw Short Fragment Buffer (SFB) at room temperature and mix by vortexing. Then spin down and keep at room temperature.
z. In a 1.5 ml Eppendorf DNA LoBind tube, mix in the following order:
   i. Library (End repair, a-tail): 60uL
   ii. Ligation Adapter (LA): 5uL
   iii. Ligation Buffer (LNB): 25uL
   iv. Salt-T4 DNA Ligase: 10uL
aa. Thoroughly mix the reaction by gently pipetting and briefly spinning down.
bb. Incubate the reaction for 10 minutes at room temperature.
cc. Resuspend the AMPure XP Beads (AXP) by vortexing.
dd. Add 50 µl of resuspended AMPure XP Beads (AXP) to the reaction and mix by flicking the tube.
ee. Incubate on a Hula mixer (rotator mixer) for 15 minutes at room temperature.
ff. Spin down the sample and pellet on a magnet. Keep the tube on the magnet, and pipette off the supernatant when clear and colourless.
gg. Wash the beads by adding 250 µl Short Fragment Buffer (SFB). Flick the beads to resuspend, spin down, then return the tube to the magnetic rack and allow the beads to pellet. Remove the supernatant using a pipette and discard.
hh. Repeat the previous step.
ii. Spin down and place the tube back on the magnet. Pipette off any residual supernatant. Allow to dry for 1 minute, but do not dry the pellet to the point of cracking.
jj. Remove the tube from the magnetic rack and resuspend the pellet in 13 µl Elution Buffer (EB). Spin down and incubate for at 37°C for 15 minutes.
kk. Pellet the beads on a magnet until the eluate is clear and colourless, for at least 1 minute.
ll. Remove and retain 13 µl of eluate containing the DNA library into a clean 1.5 ml Eppendorf DNA LoBind tube.
mm. If desired, quantify eluate using the Qubit™ dsDNA HS Quantification Assay
   i. NOTE: DNA concentration will be extremely low or too low to quantify.
nn. Store on ice until sequencing.

*Step 4: Prepare Auxiliary DNA Library*

1. Carrier DNA does not need to be the same as the input DNA for the Cas12a enrichment library. If carrier DNA is not the same as the input DNA for the Cas12a enrichment library, we recommend preparing the DNA following the manufacturer's protocol for the Native Barcoding Kit 24 V14 (Oxford Nanopore Technologies).
2. Example:
   a. Thaw the AMPure XP Beads (AXP) and mix by vortexing. Keep the beads at room temperature
   b. Prepare the NEBNext FFPE DNA Repair Mix and NEBNext Ultra II End Repair / dA-tailing Module reagents in accordance with manufacturer's instructions, and place on ice.
   c. Combine the following in a PCR tube:

| Reagent | Volume (uL) |
|---|---|
| Auxiliary gDNA | 1ug |
| NEBNext FFPE DNA Repair Buffer | 0.875 |
| Ultra II End-prep Reaction Buffer | 0.875 |
| Ultra II End-prep Enzyme Mix | 0.75 |
| NEBNext FFPE DNA Repair Mix | 0.5 |
| Water | to 15uL |

      i. Between each addition, pipette mix 10-20 times.
   d. Ensure the components are thoroughly mixed by pipetting and spin down in a centrifuge.
   e. Using a thermal cycler, incubate at 20°C for 5 minutes and 65°C for 5 minutes.
   f. Transfer the sample to a 1.5mL tube
   g. Resuspend the AMPure XP beads (AXP) by vortexing.
   h. Add 15uL of Ampure XP beads to the mix. Flick the tube to mix.
   i. Incubate on the Hula at RT for 5 minutes

j. Prepare 1mL of 80% ethanol
k. Spin down the samples and pellet the beads on a magnet until the eluate is clear and colourless. Keep the tubes on the magnet and pipette off the supernatant.
l. Keep the tube on the magnet and wash the beads with 200 µl of freshly prepared 80% ethanol without disturbing the pellet. Remove the ethanol using a pipette and discard.
m. Repeat the previous step.
n. Briefly spin down and place the tubes back on the magnet for the beads to pellet. Pipette off any residual ethanol. Allow to dry for 30 seconds, but do not dry the pellets to the point of cracking.
o. Remove the tubes from the magnetic rack and resuspend the pellet in 10 µl nuclease-free water. Spin down and incubate for 2 minutes at room temperature
p. Pellet the beads on a magnet until the eluate is clear and colourless.
q. Remove and retain 10 µl of eluate into a clean 1.5 ml Eppendorf DNA LoBind tube.
r. Quantify eluate using the Qubit™ dsDNA HS Quantification Assay
s. Prepare the NEB Blunt/TA Ligase Master Mix according to the manufacturer's instructions, and place on ice
t. Thaw the EDTA at room temperature and mix by vortexing. Then spin down and place on ice.
u. Thaw the Short Fragment Buffer (SFB) at room temperature and mix by vortexing. Place on ice.
v. Thaw the Native Barcodes at room temperature. Briefly spin down, individually mix the barcodes required for your number of samples by pipetting, and place them on ice.
w. In clean a 0.2 ml PCR-tube, add the reagents in the following order per well:

| Reagent | Volume (uL) |
|---|---|
| End-prepped DNA | 7.5 |
| Barcode 36 | 2.5 |
| Blunt/TA Ligase Master Mix | 10 |
| Total | 20 |

x. Thoroughly mix the reaction by gently pipetting and briefly spinning down.
y. Incubate for 20 minutes at room temperature
z. Add 4 µl EDTA (blue cap) to the tube and mix thoroughly by pipetting and spin down briefly.
aa. Transfer to a 1.5mL tube
bb. Resuspend the AMPure XP Beads (AXP) by vortexing.
cc. Add 10uL Ampure XP beads to the tube
dd. Incubate on a Hula mixer (rotator mixer) for 10 minutes at room temperature
ee. Spin down the sample and pellet on a magnet for 5 minutes. Keep the tube on the magnetic rack until the eluate is clear and colourless, and pipette off the supernatant.
ff. Wash the beads with 700 µl of Short Fragment Buffer (SFB). Flick the beads to resuspend, spin down, then return the sample to the magnetic rack and allow the beads to pellet. Remove the buffer using a pipette and discard.
gg. Repeat the previous step.
hh. Spin down and place the tube back on the magnetic rack. Pipette off any residual buffer.
ii. Remove the tube from the magnetic rack and resuspend the pellet in 35 µl nuclease-free water by gently flicking.
jj. Incubate for 10 minutes at 37°C. Every 2 minutes, agitate the sample by gently flicking for 10 seconds to encourage DNA elution.
kk. Pellet the beads on a magnetic rack until the eluate is clear and colourless.
ll. Remove and retain 35 µl of eluate into a clean 1.5 ml Eppendorf DNA LoBind tube.
mm. Spin down the Native Adapter (NA) and Quick T4 DNA Ligase, pipette mix and place on ice.
nn. Thaw the Elution Buffer (EB) at room temperature and mix by vortexing. Then spin down and place on ice.
oo. Thaw Short Fragment Buffer (SFB) at room temperature and mix by vortexing. Then spin down and keep at room temperature
pp. In a 1.5 ml Eppendorf LoBind tube, mix in the following order:

| Reagent | Volume (uL) |
|---|---|
| Barcoded Sample | 30 |
| Native Adapter (NA) | 5 |
| NEBNext Quick Ligation Reaction Buffer (5X) | 10 |

| | |
|---|---|
| Salt T4 DNA Ligase | 5 |
| Total | 50 |

qq. Thoroughly mix the reaction by gently pipetting and briefly spinning down.
rr. Incubate the reaction for 20 minutes at room temperature
ss. Resuspend the AMPure XP Beads (AXP) by vortexing.
tt. Add 20 µl of resuspended AMPure XP Beads (AXP) to the reaction and mix by pipetting.
uu. Incubate on a Hula mixer (rotator mixer) for 10 minutes at room temperature
vv. Spin down the sample and pellet on the magnetic rack. Keep the tube on the magnet and pipette off the supernatant.
ww. Wash the beads by adding 125 µl Short Fragment Buffer (SFB). Flick the beads to resuspend, spin down, then return the tube to the magnetic rack and allow the beads to pellet. Remove the supernatant using a pipette and discard.
xx. Repeat the previous step.
yy. Spin down and place the tube back on the magnet. Pipette off any residual supernatant.
zz. Remove the tube from the magnetic rack and resuspend pellet in 15 µl Elution Buffer (EB).
aaa. Spin down and incubate for 10 minutes at 37°C. Every 2 minutes, agitate the sample by gently flicking for 10 seconds to encourage DNA elution.
bbb. Pellet the beads on a magnet until the eluate is clear and colourless, for at least 1 minute.
ccc. Remove and retain 15 µl of eluate containing the DNA library into a clean 1.5 ml Eppendorf DNA LoBind tube.
ddd. Quantify library using the Qubit™ dsDNA HS Quantification Assay
eee. Place on ice until sequencing

*Step 5: Sequence*

1. Perform hardware check
2. Open the MinION or GridION device lid and slide the flow cell under the clip. Press down firmly on the priming port cover to ensure correct thermal and electrical contact.
3. Perform flow cell check
4. Thaw the Sequencing Buffer (SB), Library Beads (LIB) or Library Solution (LIS, if using), Flow Cell Tether (FCT) and Flow Cell Flush (FCF) at room temperature before mixing by vortexing. Then spin down and store on ice.
5. Thaw BSA
6. To prepare the flow cell priming mix with BSA, combine Flow Cell Flush (FCF) and Flow Cell Tether (FCT), as directed below. Mix by pipetting at room temperature.
    a. Flow Cell Flush (FCF): 1170 uL
    b. BSA: 5uL
    c. Flow Cell Tether (FCT): 30uL
7. Open the MinION or GridION device lid and slide the flow cell under the clip. Press down firmly on the priming port cover to ensure correct thermal and electrical contact.
8. Slide the flow cell priming port cover clockwise to open the priming port.
    a. Set a P1000 pipette to 200 µl
    b. Insert the tip into the priming port
    c. Turn the wheel until the dial shows 220-230 µl, to draw back 20-30 µl, or until you can see a small volume of buffer entering the pipette tip
9. Load 800 µl of the priming mix into the flow cell via the priming port, avoiding the introduction of air bubbles. Wait for five minutes. During this time, prepare the library for loading by following the steps below.
10. Thoroughly mix the contents of the Library Beads (LIB) by pipetting.
11. In a new 1.5 ml Eppendorf DNA LoBind tube, prepare the library for loading as follows:

| Reagent | Volume per flow cell |
|---|---|
| Sequencing Buffer (SB) | 37.5 µl |
| Library Beads (LIB) | 25.5 µl |
| Auxiliary DNA (Native Barcoded) | 1.5 |
| Cas12a Enrichment Library | 11.5 |
| Total | 75 µl |

12. Complete the flow cell priming:
    a. Gently lift the SpotON sample port cover to make the SpotON sample port accessible.

b. Load 200 µl of the priming mix into the flow cell priming port (not the SpotON sample port), avoiding the introduction of air bubbles.

13. Mix the prepared library gently by pipetting up and down just prior to loading.
14. Add 75 µl of the prepared library to the flow cell via the SpotON sample port in a dropwise fashion. Ensure each drop flows into the port before adding the next.
15. Gently replace the SpotON sample port cover, making sure the bung enters the SpotON port and close the priming port.
16. Add the light shield
17. Close the lid and start sequencing
